# IL-13 Induces a Tuft Cell-Intrinsic CD45 Checkpoint to Limit Intestinal Type 2 Immunity

**DOI:** 10.64898/2026.07.24.740376

**Authors:** C. Sochen, S. Lebon, A. Habshush-Menachem, A. Sarusi-Portuguez, V. Holiar, V. Rudenko, B. Toval, J. Liu, Y. Levin, E. Vaaknin, N. Rosenthal, N. Tiferet, I. Orr, S. Ben-Dor, R. Haffner-Krausz, R. K. Grencis, A. Munitz, D. Karo-Atar, Z. Shulman, M. Biton

## Abstract

Tuft cells initiate intestinal type 2 immunity, yet the mechanisms that restrain excessive tuft cell activation remain poorly understood. Here, we identify the receptor tyrosine phosphatase CD45 (*Ptprc*), previously considered a hematopoietic marker, as a regulator of intestinal tuft cell function. CD45 expression is restricted to a subset of tuft cells and is induced by helminth infection and IL-13. Epithelial-specific deletion of *Ptprc* activated a tuft cell inflammatory program, promoted an epithelial inflammatory state, and increased eosinophil accumulation at homeostasis. During *Heligmosomoides polygyrus bakeri* infection, CD45 deficiency enhanced ILC2 and Th2 responses and reduced parasite burden, demonstrating that epithelial CD45 limits type 2 immunity *in vivo*. Accordingly, in intestinal organoids, CD45 was dispensable for IL-13-driven tuft cell differentiation but restrained IL-13-responsive transcriptional programs. Mechanistically, CD45-deficient tuft cells exhibited altered protein abundance of STAT5 and IL17RB, implicated in tuft cell immune regulation. Together, these findings identify CD45 as a tuft cell-intrinsic regulatory checkpoint that restrains intestinal type 2 immunity through an IL-13-induced negative-feedback circuit.

## Introduction

Type 2 immunity is essential for protection against helminth infection and for tissue repair following tissue injury. However, excessive type 2 responses can promote pathological inflammation, fibrosis, and allergic disease (Lloyd and Snelgrove, 2018; Gieseck et al., 2018; Ogulur et al., 2025). This balance is particularly important in the intestine, where immune responses must rapidly eliminate parasites while maintaining tolerance toward dietary antigens, commensal microorganisms, and other luminal stimuli (Belkaid and Artis, 2013; Tang and Mullins, 2017; Loh and Tang, 2018). Understanding how protective type 2 immunity is activated while preventing excessive inflammation remains a central question in mucosal immunology.

Seminal studies demonstrated that tuft cells initiate type 2 immunity in response to helminths, protists, and microbiota-derived metabolites through production of interleukin 25 (IL-25) and other mediators (Gerbe et al., 2016; Howitt et al., 2016; von Moltke et al., 2016; Nadjsombati et al., 2018; Schneider et al., 2018). More recent work has expanded this view, revealing additional tuft cell functions in epithelial responses to bacterial and viral infection, highlighting tuft cells as central regulators of mucosal immunity (Strine and Wilen, 2022; Wilen et al., 2018; Roach et al., 2022; Xiong et al., 4 2022; Churchill et al., 2025; Fung et al., 2023).

In the small intestine (SI), tuft cells initiate a type 2 immune circuit that is critical for parasite clearance. Upon activation, tuft cells produce IL-25 that stimulates group 2 innate lymphoid cells (ILC2s) and promotes downstream Th2 responses (Gerbe et al., 2016; Howitt et al., 2016; von Moltke et al., 2016). In turn, ILC2s and Th2 cells secrete IL-13, which acts on epithelial progenitors through IL-13 receptors (IL-13R) α1 and IL-4Rα to drive differentiation toward tuft and goblet cell lineages. This feed-forward circuit amplifies type 2 immunity and facilitates parasite expulsion (Grencis, 2015; Biton et al., 2018; Xiong et al., 2022; Howitt et al., 2016; von Moltke et al., 2016; McGinty et al., 2020).

Although the mechanisms driving tuft cell-mediated immune activation have been extensively studied in the last decade, much less is known about how these responses are restrained. This question is particularly intriguing because tuft cells constitutively express IL-25 and are continuously exposed to luminal signals that can activate the tuft cell-ILC2 circuit, including microbiota-derived succinate (von Moltke et al., 2016; Nadjsombati et al., 2018; Schneider et al., 2018; Banerjee et al., 2020; Lei et al., 5 2018). Nevertheless, overt type 2 inflammation does not normally occur in the healthy intestine, suggesting the existence of regulatory mechanisms that maintain appropriate activation thresholds. Several pathways have been implicated in limiting tuft cell-driven immunity, including leukotriene signaling, IL-17rb-dependent regulation of IL-25 availability, SpiB-dependent transcriptional programs, and prostaglandin-mediated suppression of epithelial responses (Wang et al., 2025; McGinty et al., 2020; Feng et al., 2025; Oyesola et al., 2021; Xu et al., 2025). However, the upstream signaling pathways that negatively regulate tuft-cell activation thresholds still need to be elucidated.

Recent single-cell studies have revealed heterogeneity within the tuft cell lineage identifying two major tuft cell populations in mice and, more recently, four tuft cell subsets in the human small intestine (Haber et al., 2017; Billipp et al., 2024; Buissant des Amorie et al., 2025). In particular, the tuft-1 subset is enriched for a neuronal-associated gene signature, while the tuft-2 subset exhibits an immune-enriched transcriptional program, including Siglec-F and mediators of leukotriene biosynthesis (Haber et al., 2017; McGinty et al., 2020; Feng et al., 2024). Intriguingly, tuft-2 cells selectively express *Ptprc* (CD45) (Haber et al., 2017). CD45 is a receptor-type tyrosine phosphatase traditionally considered a pan-leukocyte marker and a central regulator of signaling thresholds in hematopoietic cells (Al Barashdi et al., 2021; Hermiston et al., 2003; Saunders and Johnson, 2010; Courtney et al., 2019; Irie-Sasaki et al., 2001). Through modulation of Src family kinases, JAK kinases, and other signaling intermediates, CD45 can either enhance or suppress immune activation depending on cellular context (Hermiston et al., 2003; Saunders and Johnson, 2010; Courtney et al., 2019; Irie-Sasaki et al., 2001). Notably, several signaling molecules, such as Hck from the Src family kinases and JAK1, are known to interact with CD45 and are also highly expressed in tuft cells, suggesting that CD45 may function as a regulator of tuft immune signaling. However, whether CD45 has a physiological role in non-hematopoietic cells remains largely unknown.

Here, we show that CD45 expression is enriched in a subset of intestinal tuft cells, induced by IL-13 and helminth infection, and functions to restrain tuft cell activation and downstream type 2 immune responses. Using epithelial-specific deletion of *Ptprc*, we demonstrate that CD45 limits epithelial inflammatory programs, constrains ILC2 and Th2 responses during *Heligmosomoides polygyrus bakeri* (*Hp*) infection, and regulates molecular pathways associated with tuft cell activation. Together, our findings reveal an IL-13-induced epithelial checkpoint that restrains tuft cell activation and establishes a negative feedback circuit that limits intestinal type 2 immunity.

## Results

### CD45 is selectively expressed by intestinal tuft cells and is enriched in the distal SI

To investigate the potential involvement of CD45 in tuft cell biology, we began with exploring its expression and regional distribution along the SI. Re-analysis of a published single-cell RNA-sequencing atlas of the intestinal epithelium (Haber et al., 2017) revealed that *Ptprc* expression within the epithelial compartment is restricted to tuft cells (**Fig. 1 A**). Within the tuft cell compartment, *Ptprc* expression was largely confined to the tuft-2 subset (**Fig. 1 B**).

**Figure 1.**
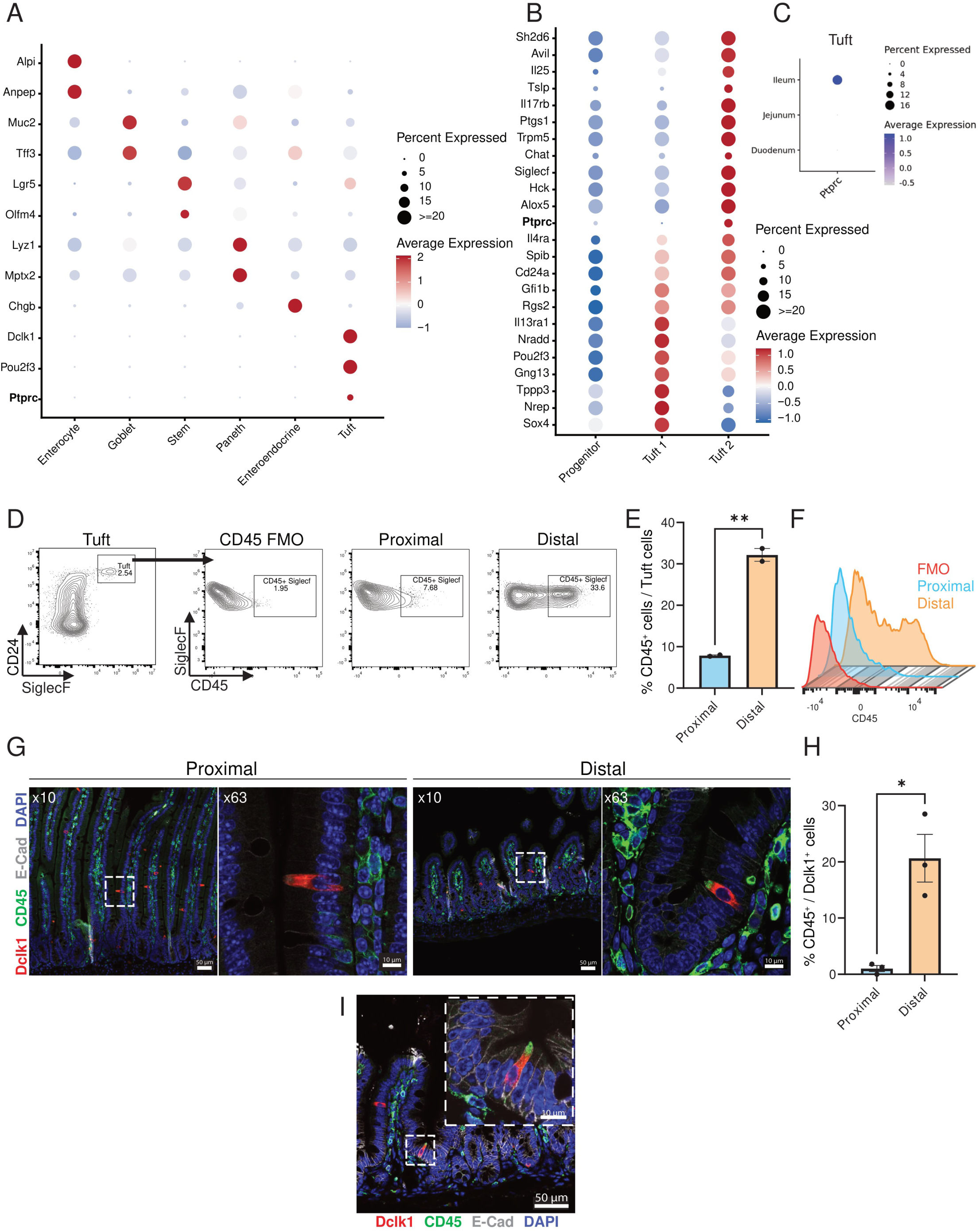
CD45-expressing tuft cells are enriched in the distal small intestine. (a-c) Re-analysis of a published single cell RNA-sequencing dataset of the mouse small intestinal epithelium (Haber et al., 2017). **(A)** Dot plot showing expression of epithelial lineage marker genes across epithelial cell populations. **(B)** Dot plot showing expression of tuft progenitor, tuft-1, and tuft-2 signature genes. **(C)** dot plot of regional expression of *Ptprc* across small intestinal epithelial populations. **(D-F)** Cytometry analysis of CD45^+^ tuft cells. **(D)** Representative flow cytometry analysis of CD45 expression in tuft cells isolated from the proximal and distal small intestine. Tuft cells were identified as live EpCAM⁺ CD45^neg-mid^ SIGLECF⁺ CD24⁺ epithelial cells. Fluorescence-minus-one (FMO) controls were used to define the CD45 gate. **(E)** Quantification of the frequency of CD45⁺ tuft cells in the proximal and distal small intestine. **(F)** Representative histograms showing CD45 expression by proximal and distal tuft cells. **(G-H)** Immunofluorescence (IF) analysis of CD45^+^ tuft cells. **(G)** Representative immunofluorescence images of proximal and distal small intestine stained for DCLK1 (red), CD45 (green), E-cadherin (gray), and DAPI (blue); scale bar, 50 μm. Insets show high-magnification images of representative tuft cells; scale bar, 10 µm. **(H)** Quantification of CD45⁺DCLK1⁺ tuft cells in the proximal or distal regions. Quantification was assessed by 10 fields of view (FOV) of 3 mice per group **(I)** Representative immunofluorescence image demonstrating apical localization of CD45 in a DCLK1⁺ tuft cell; scale bar, 50 µm. The inset shows a high-magnification view; scale bar, 10 µm. Data are shown as mean ± SEM. Statistical significance was determined using an unpaired two-tailed Student’s *t*-test. \**P* ≤ 0.05; \*\**P* ≤ 0.01.

Recent studies have highlighted the regional specialization of tuft cells, including the enrichment of Chat⁺ tuft cells in the distal SI (Billipp et al., 2024). Notably, *Chat* expression has been associated with the tuft-2 transcriptional program and distal SI identity (Billipp et al., 2024; Buissant des Amorie et al., 2025). Using Chat-GFP reporter mice, we confirmed a higher frequency of Chat⁺ tuft cells in the distal compared to proximal SI (14.73% vs. 4.97%; **Fig. S1, A-B**). Within the Chat⁺ compartment, CD45⁺ tuft cells were also more abundant distally (27.5% vs. 5.6%; **Fig. S1 C**), although CD45 and Chat expression did not fully overlap, suggesting heterogeneity within distal tuft cells. Consistent with these findings, *Ptprc* expression in publicly available datasets was largely restricted to the ileum, with minimal expression in proximal SI regions (Haber et al., 2017) (**Fig. 1 C**). We next validated this regional enrichment at the protein level by flow cytometry. Following exclusion of hematopoietic cells and gating on EpCAM⁺ epithelial cells (**Fig. S1 A**), tuft cells (CD24⁺ SiglecF⁺(McGinty et al., 2020) displayed markedly higher CD45 expression in the distal compared to proximal SI (20.1% vs. 2.26%; **Fig. 1, D-F**). Immunofluorescence (IF) analysis further confirmed that CD45 is expressed by a subset of DCLK1⁺ tuft cells, with a significantly higher frequency in the distal SI (20.65%) than in the proximal SI (1.02%) (**Fig. 1, G-H**). Importantly, CD45 expression was not detected in other epithelial cell types, indicating specificity to tuft cells within the intestinal epithelium.

Given the enrichment of CD45⁺ tuft cells in the distal SI, a region characterized by increased microbial density and diversity (Vuik et al., 2019; Delbaere et al., 2023), we next examined the subcellular localization of CD45. Strikingly, CD45 exhibited polarized apical localization in all CD45⁺ tuft cells analyzed, facing the intestinal lumen (**Fig. 1 I**), reminiscent of other tuft cell-expressed immune receptors, such as SiglecF and CD24 (McGinty et al., 2020; Billipp et al., 2024). Together, these data identify CD45 as a tuft cell-specific marker enriched in the distal SI predominantly within the tuft-2 subset. Accordingly, subsequent analyses focused on this region.

### Epithelial-specific deletion of CD45 induces a tuft cell activation program and disrupts epithelial and immune homeostasis

CD45 (*Ptprc*) is broadly expressed in hematopoietic cells (Hermiston et al., 2003), precluding the use of germline knockout models to study its epithelial function. To overcome this limitation, we generated an inducible epithelial-specific CD45 knockout model. Given the presence of multiple alternatively spliced *Ptprc* isoforms (Tchilian and Beverley, 2006), we inserted loxP sites flanking the promoter and conserved exons 1-2 using CRISPR-Cas9 (**Fig. 2 A, Methods**), and crossed these mice with Villin-CreER^T2^ animals to generate inducible and specific CD45 epithelial knock-out (CD45^ΔIEC^) mice. As CD45 expression within the intestinal epithelium is restricted to tuft cells (**Fig. 1 A**), this model enables selective interrogation of CD45 function in tuft cells *in vivo*. We then used Tamoxifen administration for 10 days in adult mice to efficiently induce CD45 deletion in the intestinal epithelium (**Fig. 2 B**). Analysis of FACS-isolated tuft cells (CD24^hi^FSC-A^low^**, Fig. S2 A**) demonstrated loss of *Ptprc* expression in CD45^ΔIEC^ mice, without affecting the expression of the tuft lineage-defining transcription factor *Pou2f3* (**Fig. 2 C**). At the protein level, flow cytometry and IF analysis confirmed efficient ablation of CD45 in tuft cells, with a 3.2-fold reduction of DCLK1⁺CD45⁺ cells in CD45^ΔIEC^ mice compared to controls (**Fig. 2, D-G, Fig. S2 B**). CD45 deletion was specific to the epithelial compartment, as CD45 expression was unchanged in lamina propria and Peyer’s patches immune cells in both flow cytometry and IF analysis. (**Fig. 2 F, Fig. S2, C-D**). Histological analysis revealed no overt changes in tissue architecture or inflammation, indicating that CD45 loss does not grossly disrupt epithelial integrity at baseline (**Fig. 2 I**). In addition, we did not observe changes in cell proliferation (**Fig. S2 F**). Notably, tuft cell abundance remained similar in CD45 deficient mice, as observed by flow cytometry and IF analysis (**Fig. 2 H, Fig. S2 E**).

**Figure 2.**
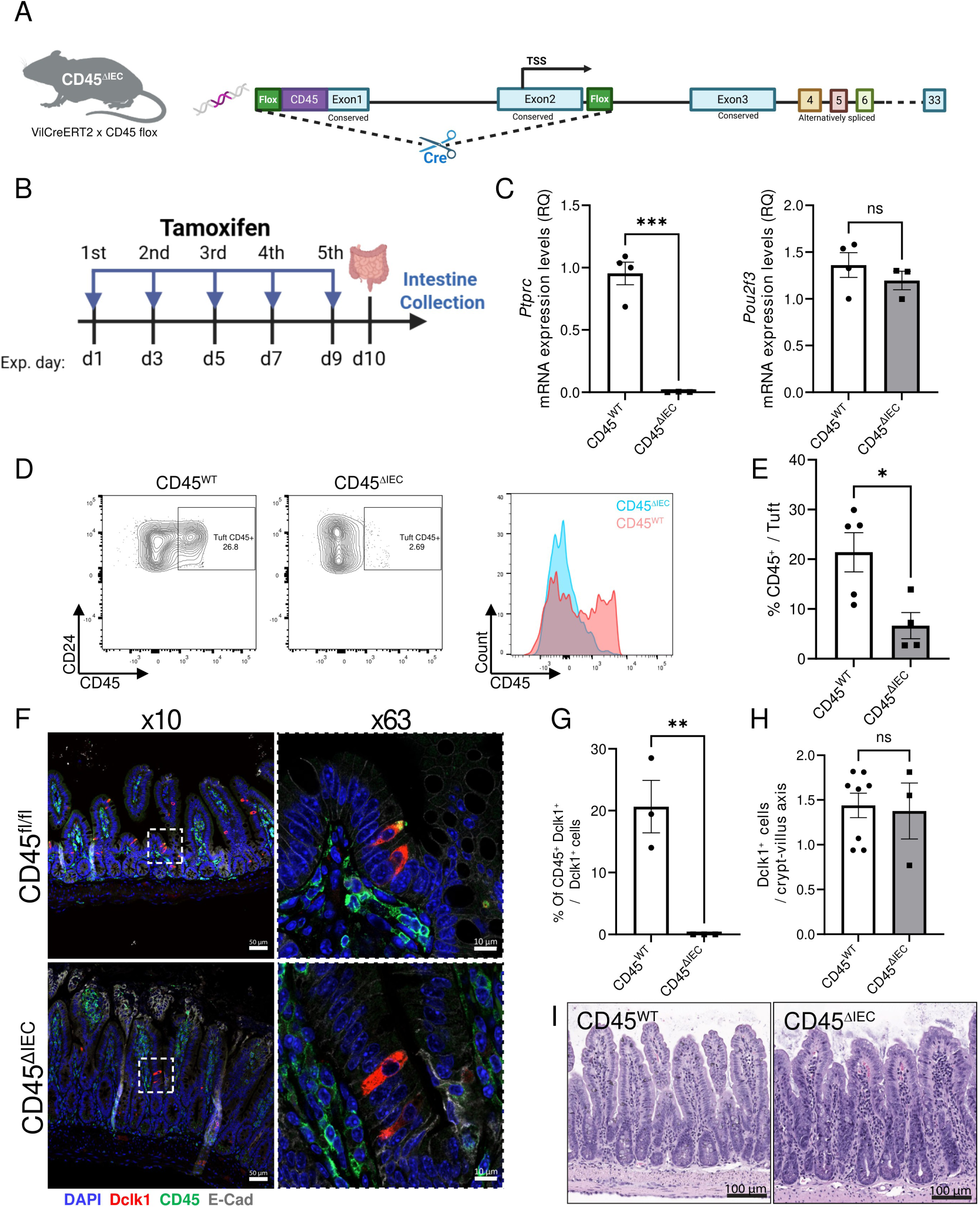
Generation and validation of an inducible epithelial-specific CD45 knockout mouse model. **(A)** Schematic of the inducible epithelial-specific *Ptprc* knockout mouse model (CD45^ΔIEC^), generated by crossing *Ptprc*^fl/fl^ mice with Villin-CreERT2 mice. LoxP sites flank the promoter and exons 1-2 of *Ptprc*. Schematics were created with BioRender. **(B)** Experimental timeline showing tamoxifen administration and tissue collection. **(C)** Quantitative PCR (qPCR) analysis of *Ptprc* (left) and *Pou2f3* (right) expression in FACS-isolated tuft cells from CD45^ΔIEC^ and CD45^WT^ control mice. **(D-E)** Validation of CD45 depletion in tuft cells by cytometry analysis. **(D)** Representative flow cytometry analysis of CD45 expression in tuft cells from CD45^ΔIEC^ and CD45^WT^ mice. Tuft cells were identified as live EpCAM⁺ CD45^+^ CD24⁺ FSC-A^low^ epithelial cells. Representative histograms show CD45 expression. **(E)** Quantification of the frequency of CD45⁺ tuft cells determined by flow cytometry. n = 5 mice per group. **(F-H)** Validation of CD45 depletion by immunofluorescence analysis. **(F)** Representative immunofluorescence images of distal small intestine sections stained for DCLK1 (red), CD45 (green), E-cadherin (gray), and DAPI (blue). **(G)** Quantification of CD45⁺DCLK1⁺ tuft cells from immunofluorescence images. Quantification was performed using 10 fields of view (FOVs) from 3 mice per group. **(H)** Quantification of DCLK1⁺ tuft cells along the crypt-villus axis. Quantification was performed using 10 fields of view (FOVs) from 8 mice in the CD45^WT^ group and 3 mice in the CD45^ΔIEC^ group. **(I)** Representative hematoxylin and eosin (H&E)-stained sections of distal small intestine from CD45^ΔIEC^ and CD45^WT^ mice. Scale bar, 100 µm. Data are shown as mean ± SEM. Statistical significance was determined using an unpaired two-tailed Student’s *t*-test. \**P* ≤ 0.05; \*\**P* ≤ 0.01, \*\*\**P* ≤ 0.001; ns, not significant.

To define the functional consequences of tuft CD45 loss, we performed bulk RNA sequencing on FACS-isolated tuft cells (EpCAM⁺CD24^hi^FSC-A^low^) (Haber et al., 2017) and total epithelial cells (EpCAM⁺) from CD45^ΔIEC^ and control mice (**Fig. 3, A-C**, **Table S1, Methods**). As expected, in tuft cells *Ptprc* was the most significantly downregulated gene, confirming efficient deletion (**Fig. 3 A**). Notably, CD45-deficient tuft cells exhibited upregulation of genes associated with inflammatory and stress signaling pathways, including components of the TNF and NF-κB pathways (*Wdfy1*, *Chuk*, *Rac1*, *Zfand5*, *Traf4*), as well as regulators of oxidative stress and innate signaling such as *Gclm* and *Selenoo*. Increased expression of *Gata4*, a transcription factor linked to inflammation and antimicrobial defense in the small intestine(Haveri et al., 2009; Earley et al., 2023), further supported the acquisition of an immune-responsive state. Gene ontology (GO) analysis revealed enrichment of pathways associated with cellular stress adaptation, including autophagy, mitochondrial function, and endoplasmic reticulum stress, alongside activation of inflammatory signaling modules such as NF-κB, Toll-like receptor, TNF, chemokine, and IL-17 pathways (**Fig. 3 B, Fig. S3 A**). These changes were reflected in the differential expression of key regulatory and signaling genes (e.g., *Chuk*, *Traf4*, *Mapk1*, *Rac1*, *Cebpd*, *Wdfy1*), as well as genes involved in intracellular trafficking and mitochondrial quality control (e.g., *Lamp1*, *Pink1*) (**Fig. 3, A** and **C**, and **Fig. S3 A**).

**Figure 3.**
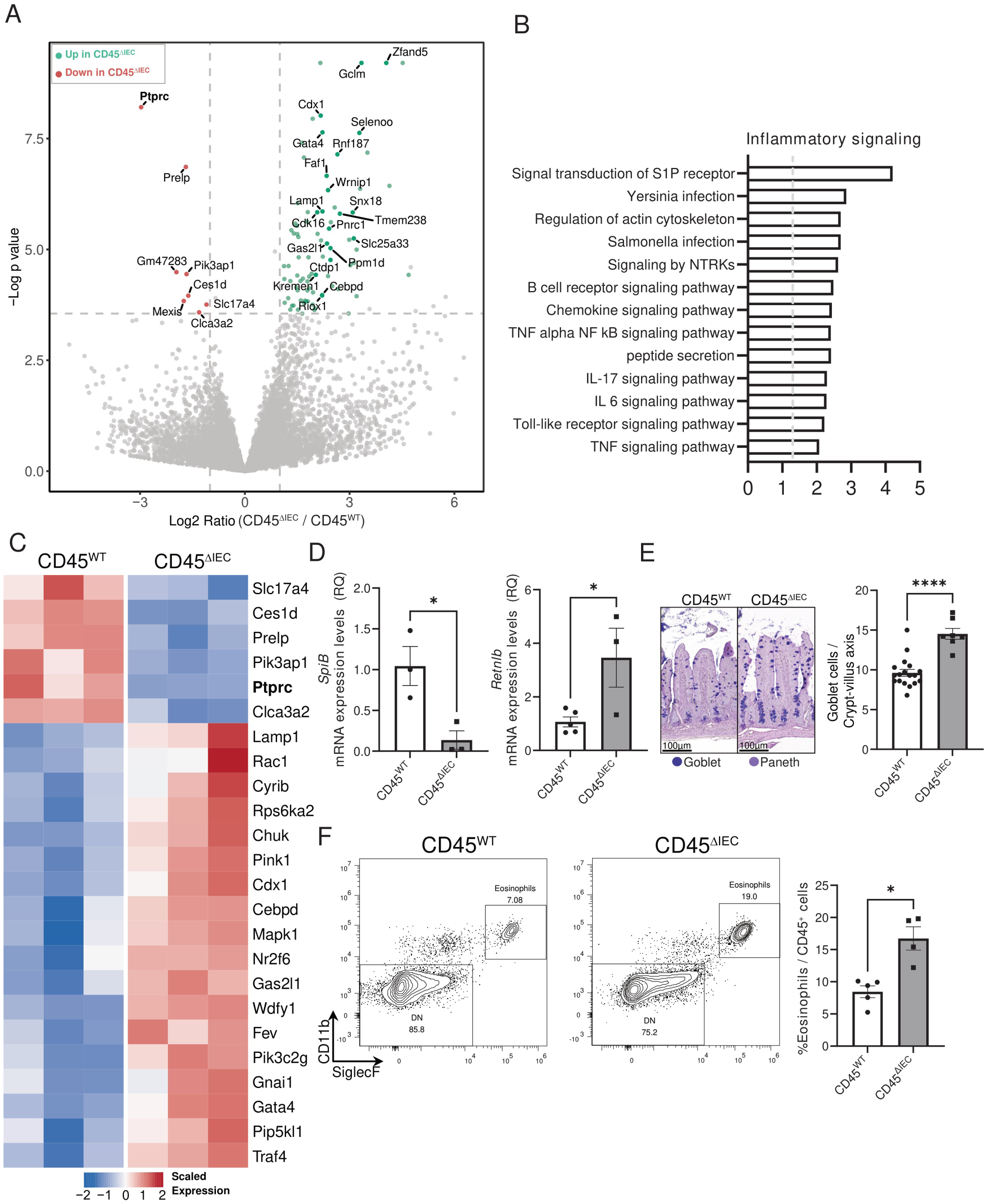
Epithelial CD45 deficiency induces a tuft-cell inflammatory program and disrupts epithelial and immune homeostasis. **(A-C)** Transcriptomic analysis of FACS-isolated tuft cells and intestinal epithelial cells (IECs) from CD45^ΔIEC^ and CD45^WT^ mice. Tuft cells were isolated as CD24⁺ FSC-A^low^ epithelial cells and IECs as EpCAM⁺ cells. **(A)** Volcano plot showing differentially expressed genes in CD45^ΔIEC^ versus CD45^WT^ tuft cells. Upregulated genes are shown in green, downregulated genes in red, and unchanged genes in gray. Selected genes are indicated. Vertical dashed lines indicate the differential expression threshold (|log₂ fold change| > 1), and the horizontal dashed line indicates the significance threshold (FDR < 0.05). **(B)** Gene Ontology enrichment analysis of upregulated genes in CD45^ΔIEC^ tuft cells, highlighting inflammatory signaling pathways. **(C)** Heatmap showing representative differentially expressed genes in tuft cells from CD45^ΔIEC^ and CD45^WT^ mice. **(D)** qPCR analysis of selected epithelial genes in EpCAM⁺ cells isolated from CD45^ΔIEC^ and CD45^WT^ mice, *n* = 3-5 mice per group. **(E)** Representative Alcian blue-periodic acid-Schiff (AB-PAS) staining of distal small intestine sections (left) and quantification of goblet cells along the crypt-villus axis (right). Scale bar, 100μm. **(F)** Representative flow cytometry analysis (left) and quantification (right) of eosinophils in the lamina propria of CD45^ΔIEC^ and CD45^WT^ mice, *n* = 4-5 mice per group. Data are shown as mean ± SEM. Statistical significance was determined using an unpaired two-tailed Student’s *t*-test. \**P* ≤ 0.05; \*\*\*\**P* ≤ 0.0001; ns, not significant.

Importantly, depletion of CD45 in tuft cells was accompanied by extensive transcriptional remodeling of the broader epithelial compartment. Bulk EpCAM⁺ RNA-seq identified 1,304 upregulated and 169 downregulated genes in CD45^ΔIEC^ mice relative to controls (**Fig. S3 B**). Upregulated genes included inflammatory mediators (*Il18*, *Il18bp*, *Il21r*, *Ccl5*, *Ccl25*, *Areg*) and antimicrobial effectors (*Reg3a*, *Reg3b*, *Reg3g*, *Retnlb*, *Saa1*, *Saa3*, *Sprr2a3*). In contrast, genes associated with epithelial differentiation and secretory function, including *Atoh1*, *Sox9*, *Lgr5*, *Muc2*, *Il17re*, *Spib*, and *Kit*, were downregulated. GO analysis confirmed enrichment of immune and inflammatory processes, alongside reduced representation of epithelial differentiation, metabolic, and secretory programs (**Fig. S3 C**). These findings were validated by qPCR, which confirmed decreased *Spib* expression, a microfold (M) and tuft cells transcription factor, whose downregulation was recently shown to promote tuft cell-mediated type 2 immunity activation (Wang et al., 2025). In parallel, a significant increase in the expression of *Retnlb,* a known type 2 induced gene in goblet cells, which play a role in parasite immunity, was validated in CD45^ΔIEC^ mice (Artis et al., 2004) (**Fig. 3 D**). Consistent with the elevated expression of *Retnlb*, Alcian Blue-Periodic Acid-Schiff (AB-PAS, **Methods)** stain indicated increased goblet cell numbers in CD45^ΔIEC^ mice (**Fig. 3 E**). Given that tuft cells comprise a minor fraction of the epithelium, and CD45⁺ tuft cells represent only a subset of this population, the magnitude of epithelial transcriptional changes suggests that CD45-expressing tuft cells act as regulators of epithelial homeostasis.

Finally, we assessed whether epithelial CD45 deletion affects the immune compartment, in particular type 2 immunity. Given the enrichment of eosinophil-associated pathways and chemokines in the GO analysis, and the established role of tuft cells in type 2 immunity, we performed flow cytometric analysis of lamina propria immune populations (**Fig. S3, D-E**). CD45^ΔIEC^ mice exhibited an approximately 2-fold increase in eosinophil frequency (**Fig. 3 F**), while ILC2 and Th2 abundance remained unchanged (**Fig. S3 F**). In addition, no change in Th1 abundance was observed (**Fig. S3 F**). Notably, regulatory T cells (Tregs) were increased, whereas Th17 cells were reduced (**Fig. S 3 G**), indicating a shift in gut-specialized CD4^+^ T cell immune balance.

Together, the induction of epithelial *Retnlb* (RELMβ) and *Areg*, two mucosal mediators associated with type 2-linked tissue responses (Herbert et al., 2009; Stephen-Victor et al., 2025; Zaiss et al., 2006, 2015; Morimoto et al., 2018; Huang et al., 2024), with eosinophil expansion in CD45^ΔIEC^ mice, supports the idea that CD45-depleted tuft cells promote a mild type 2-skewed inflammatory landscape. However, this response was not accompanied by increased ILC2 or Th2 cell abundance in the distal small intestines.

### CD45 deletion during *Heligmosomoides polygyrus* infection exacerbates type 2 immune responses and reduces infection burden

To further investigate the possible role of tuft-CD45 in regulating type 2 immune responses, we set out to explore epithelial CD45 deletion in the context of a well-known type 2 immunity inducer, *Heligmosomoides polygyrus bakeri* (*Hp*) infection (Johnston et al., 2015; Reynolds et al., 2012). To induce CD45 deletion, we injected CD45^fl/fl^ or CD45^ΔIEC^ mice with tamoxifen for 10 days, then infected mice with 200 *Hp* Larvae and collected the intestine 10 days post-infection (Johnston et al., 2015) (**Fig. 4 A** and **Methods**). In infections induced by oral gavage in WT mice, *Hp* parasites are found in the small intestine, particularly the duodenum, throughout all stages of their life cycle (Reynolds et al., 2012). We first confirmed successful infection by observing adult *Hp* worms residing in the duodenal SI (**Fig. S4 A**). Importantly, CD45 was successfully deleted according to flow cytometry analysis, and qPCR **(Fig. 4A, Fig. S4 B)**. One of the known consequences of *Hp* infection is expansion of the tuft cell population in the proximal SI (Gerbe et al., 2016; Howitt et al., 2016; von Moltke et al., 2016; Biton et al., 2018). Therefore, we examined whether tuft cells also expand in the distal SI, our region of interest, by flow cytometry and IF tissue staining (**Fig. S4, C-D**). We observed no significant expansion in distal SI tuft cells following Hp infection in both CD45^ΔIEC^ and control CD45^fl/fl^ mice.

**Figure 4.**
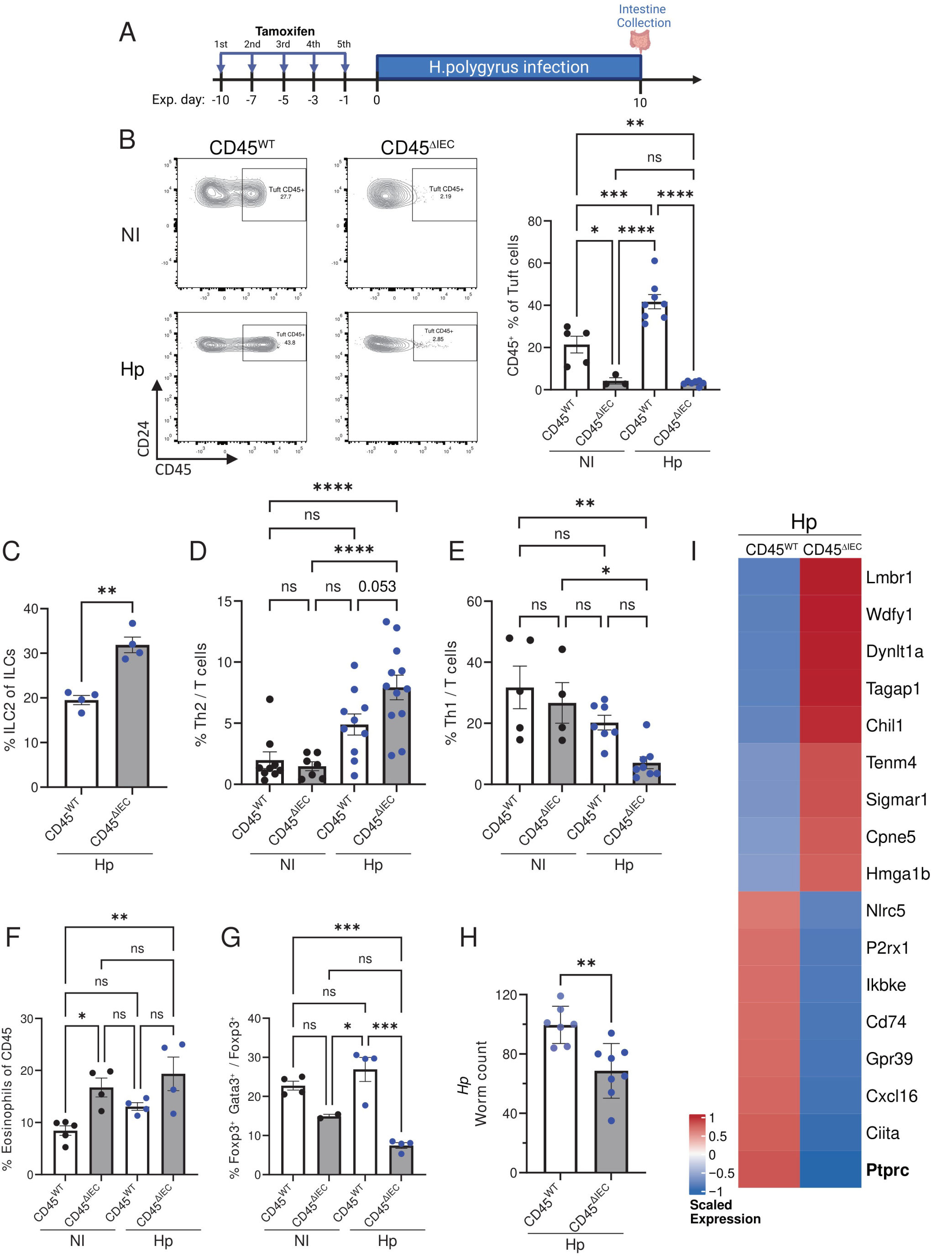
Epithelial CD45 deficiency enhances type 2 immunity and reduces parasite burden during *Heligmosomoides polygyrus* infection. **(A)** Experimental design. CD45^ΔIEC^ and CD45^WT^ mice were treated with tamoxifen, infected with *Heligmosomoides polygyrus bakeri* (*Hp*), and analyzed 10 days after infection. Schematic created with BioRender. **(B)** Representative flow cytometry plots and quantification of CD45⁺ tuft cells from the distal small intestine of noninfected (NI) and *Hp*-infected CD45^ΔIEC^ and CD45^WT^ mice, n = 3-8 mice per group. **(C)** Flow cytometry quantification of lamina propria ILC2 frequencies in *Hp*-infected CD45^ΔIEC^ and CD45^WT^ mice, n = 4 mice per group. **(D-G)** Flow cytometric analysis of lamina propria immune populations from noninfected and *Hp*-infected CD45^ΔIEC^ and CD45^WT^ mice. Quantification of **(D)** Th2 cells (n = 7-12), **(E)** Th1 cells (n = 4-8), **(F)** eosinophils (n = 4-5), and **(G)** Foxp3⁺GATA3⁺ regulatory T cells (Tregs, n = 2-5). **(H)** Quantification of adult *Hp* worms recovered from the proximal small intestine. n = 7-8 mice per group. **(I)** Heatmap showing selected differentially expressed genes, associated with the defense transcriptional program of tuft cells isolated from *Hp*-infected CD45^ΔIEC^ (n = 5) and CD45^WT^ (n = 4) mice and sequenced by bulk RNA sequencing. Data are shown as mean ± SEM. Statistical significance was determined using unpaired two-tailed Student’s *t* test (**C**, **H**) or one-way ANOVA followed by Tukey’s multiple-comparison test (**B**, **D-G**, **I**). \**P* ≤ 0.05; \*\**P* ≤ 0.01; \*\*\**P* ≤ 0.001, \*\*\*\**P* ≤ 0.0001; ns, not significant.

We next examined CD45 expression in tuft cells during infection. Flow cytometry analysis revealed an approximately two-fold increase in CD45 expression in tuft cells from WT-infected mice, from ∼21% in non-infected (NI) mice to 42% in Hp-infected mice (**Fig. 4 B**). This increase in *Ptprc* expression was also observed by qPCR analysis of sorted tuft cells (**Fig. S4 B**). To further verify the induction of CD45 expression in *Hp* infection in the distal SI, we examined scRNA-seq data from Haber et al. (Haber et al., 2017). We observed a minor increase on day 3 of infection, with a more substantial increase on day 10, in agreement with our observations (**Fig. S4 E).** Finally, we did not observe differences in tissue architecture or damage as a result of CD45 deletion in infected mice **(Fig. S4 A)**. In addition, no changes in tuft cell proliferation were noticed as measured by nuclear staining of mKi67, in *Hp*-infected control or CD45-deficient mice (**Fig. S4 F**).

Studies have shown that tuft cells initiate and recruit ILC2 and Th2 cells upon *Hp* infection (McGinty et al., 2020; Schneider et al., 2018; Howitt et al., 2016; von Moltke et al., 2016); thus, we characterized the corresponding populations in CD45^ΔIEC^ or control CD45^fl/fl^ mice. Infected CD45^ΔIEC^ mice exhibit a greater expansion of ILC2 and Th2 cells compared with control mice (**Fig. 4, C-D**). Consistent with these results, the Th1 population was inversely reduced in infected knockouts **(Fig. 4 E)**. Tregs and Th17 cells which shifted their proportions in CD45 deficient mice under homeostasis (**Fig. S3 G**), exhibited similar changes under *Hp* infection **(Fig. S4 G**). Similarly, eosinophils increased in CD45 deficient mice, regardless of infection (**Fig. 4 F)**. We then examined the Gata3^+^ Treg population, which is associated with restraining type 2 immunity (Wang et al., 2011; Ding et al., 2022). We observed a significant reduction in Foxp3^+^ Gata3^+^ Tregs in CD45-deficient mice following infection, consistent with the overall increase in type 2 immunity **(Fig. 4 G)**.

Due to the stronger type 2 immune response in CD45^ΔIEC^ mice, we asked whether this affects *Hp* infection burden. Quantification of adult *Hp* worms in the duodenal SI tissue reveals a significant decrease in worm burden in CD45-deficient mice at day 10 **(Fig. 4 H**). These results demonstrate that in the absence of CD45 expression in tuft cells, type 2 immune responses are amplified and are associated with improved control of *Hp* infection.

To determine whether the amplified immune phenotype in infected CD45^ΔIEC^ mice was accompanied by tuft-intrinsic transcriptional changes, we performed bulk RNA sequencing of tuft cells isolated from CD45^ΔIEC^ mice and CD45^fl/fl^ controls that were *Hp*-infected (*Hp*) (**Fig. 4 I and Table S2**). Infected CD45^ΔIEC^ tuft-cell transcriptome revealed a coordinated shift, with upregulation of a discrete epithelial defense-associated program and downregulation of genes linked to antigen presentation, cytokine-mediated signaling, and immune activation (**Fig. 4 I** and **Fig. S4 H**). In parallel, infected CD45^ΔIEC^ tuft cells upregulated a coordinated transcriptional program associated with innate defense, inflammatory signaling, and epithelial interactions with environmental cues. Representative genes within this program included *Wdfy1*, which has been implicated in TLR-dependent inflammatory signaling and type I interferon responses, *Tagap*, which has been linked to antimicrobial effector pathways, and *Chil1*, which has been associated with host–microbiota interactions (Yang et al., 2020; Chen et al., 2020, 2024; He et al., 2022). Together, these findings reveal a distinct tuft cell activation state in the absence of CD45, linking enhanced type 2 immunity and reduced parasite burden to an activated tuft cell transcriptional program.

As local type 2 immunity was elevated, we next asked whether these changes extended to the draining mesenteric lymph nodes (MLNs) (**Fig. S5**). During *Hp* infection, MLNs are known to exhibit expansion of plasma cells, germinal center (GC) B cells, and T follicular helper (Tfh) cells, which support their expansion (Harris and Gause, 2011; Reynolds et al., 2012). In addition, B cells undergo class switching mainly to IgG1 and IgE, although IgE is not essential for host protection in *Hp* infection (Harris and Gause, 2011). Under normal conditions, we observed these populations were kept at basal levels, regardless of mouse genotype (**Fig. S5, A-C**). As expected, in CD45^fl/fl^ mice during *Hp* infection (Day 10) we observed a significant increase in the frequencies of B cells, GC, and plasma B cells (**Fig. S5, A-C).** In contrast, in epithelial CD45-deficient mice, plasma and GC cells remained at basal levels despite infection (**Fig. S5, B-C)**. We next examined the B cell response by analyzing immunoglobulin (Ig) isotype frequencies in the MLNs **(Fig. S5 D)**. We observed no significant differences in IgG1 or IgG2c isotypes in the MLNs of epithelial CD45-deficient mice, whether under normal conditions or during *Hp* infection. In addition, IgM B cells were similarly increased in both genotypes under infection. In contrast, IgA^+^ GC B cells were significantly increased in infected mice deficient for epithelial CD45, suggesting a stronger barrier defense response program in these mice. Together, epithelial CD45 deletion was associated with both enhanced local type 2 effector responses in the intestine, alongside a markedly altered MLN B cell response to *Hp* infection.

### CD45 in tuft cells restricts tuft cell activation in response to IL-13, but not tuft cell differentiation

We next aimed to determine the functional connection between enhanced type 2 immunity and the transcriptional changes induced by CD45 epithelial deletion. A key player of type 2 immunity in the *Hp*-tuft-ILC2 circuit is the IL-13 cytokine (Howitt et al., 2016; von Moltke et al., 2016; Gerbe et al., 2016). IL-13 is secreted from ILC2 and Th2 cells and is a strong driver of secondary epithelial transcriptional changes upon helminth infection, including the expansion of tuft and goblet cells from Lgr5^+^ stem cells (Biton et al., 2018).

Hence, we set out to specifically investigate whether CD45 expression affects tuft cell differentiation in the presence of IL-13. We thereby cultured distal small intestine organoids from CD45^ΔIEC^ mice or matched controls, stimulating them at the passaging stage with IL-13 for 72 hours **(Fig. 5 A, Fig. S6 A**, **Methods**). We first examined the expression of known tuft markers via qPCR. The expression of *Dclk1*, *Pou2f3*, and *IL-25* significantly increased following IL-13 stimulation and did not vary in levels between control and CD45^ΔIEC^ organoids (**Fig. 5 B)**. At the population level, IF staining revealed significant yet comparable expansion of DCLK1^+^ tuft cells in both IL-13-stimulated groups compared to non-stimulated controls (**Fig. 5, C-D)**. Furthermore, RNA-seq analysis of these organoids reveals a similar tuft cell signature between IL-13-stimulated groups, except for *Ptprc*, which was substantially lower in the CD45^ΔIEC^ group (**Fig. S6, B-C** and **Table S3)**. Thus, lack of epithelial-specific CD45 expression does not alter organoid differentiation capacity to tuft cells upon IL-13 stimulation.

**Figure 5.**
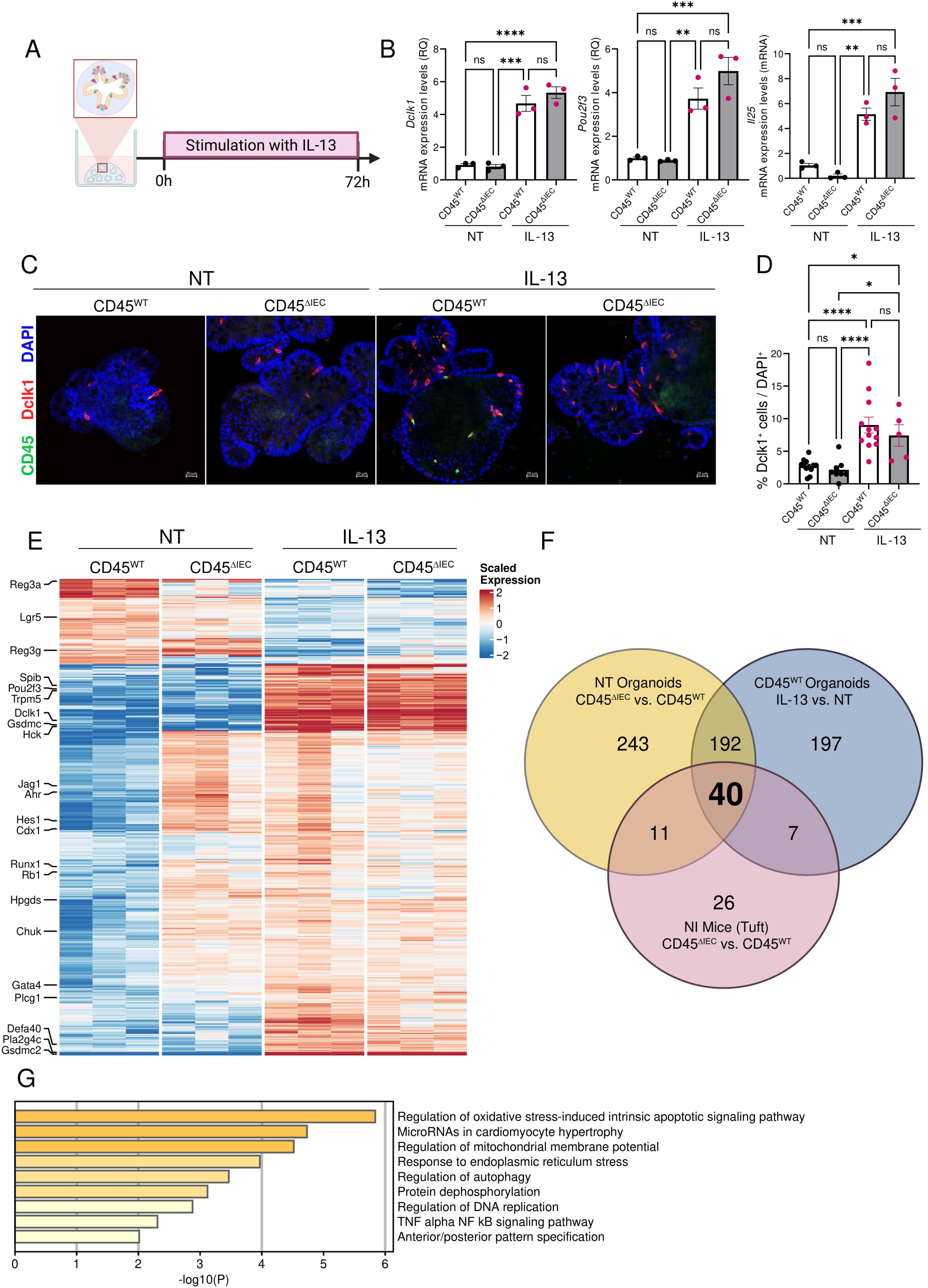
CD45 restrains IL-13-induced tuft cell activation but is dispensable for tuft cell differentiation. **(A)** Experimental design. Distal small intestinal organoids derived from CD45^ΔIEC^ and CD45^WT^mice were cultured in the presence or absence of IL-13 for 72 h before analysis. Schematic created with BioRender. **(B)** qPCR analysis of tuft-cell marker genes (*Dclk1*, *Pou2f3*, and *Il25*) in non-treated (NT) and IL-13-treated organoids, n = 3 organoid well replicates per group. **(C)** Representative immunofluorescence images of non-treated (NT) and IL-13-treated organoids stained for DCLK1 (red), CD45 (green), E-cadherin (gray), and DAPI (blue). Scale bar, 20μm. **(D)** Quantification of DCLK1⁺ tuft cells as the percentage of DCLK1⁺ cells per DAPI⁺ cells in individual organoids. Each dot represents the average percentage per one organoid, n = 5-12 organoids per condition were quantified. **(E)** Heatmap of differentially expressed genes identified by bulk RNA sequencing of non-treated (NT) and IL-13-treated CD45^ΔIEC^ and CD45^WT^ organoids, *n* = 3 organoid well replicates per group. **(F)** Venn diagram showing the overlap between upregulated genes in non-treated CD45^ΔIEC^ versus CD45^WT^ organoids, CD45^ΔIEC^ versus CD45^WT^ tuft cells *in vivo*, and IL-13-treated versus untreated CD45^WT^ organoids. Forty genes were upregulated and shared across all three datasets. **(G)** Gene Ontology enrichment analysis of the 40-gene shared transcriptional program identified in panel **F**. Data are shown as mean ± SEM. Statistical significance was determined using one-way ANOVA followed by Tukey’s multiple-comparison test (**b**, **d**). \**P* ≤ 0.05; \*\**P* ≤ 0.01; \*\*\**P* ≤ 0.001; \*\*\*\**P* ≤ 0.0001; ns, not significant.

Several studies suggest that tuft-1 and tuft-2 cells represent sequential maturation states, with tuft-1 cells serving as precursors to tuft-2 cells, which are associated with immune responses (Haber et al., 2017; Manco et al., 2021; Buissant des Amorie et al., 2025). As we observed a skew toward type 2 responses in CD45^ΔIEC^ mice, we sought to determine whether this phenomenon is mediated by a higher abundance of tuft-2 cells. Therefore, we examined the tuft-1 and tuft-2 signature scores in our organoid RNA-seq data (**Fig. S6 D, Table S3**). Although CD45 is a hallmark of tuft-2 cells, the scores of both tuft-1 and tuft-2 signatures remain unchanged in *Ptprc*-deleted organoids. Thus, CD45 expression does not affect IL-13-induced tuft cell differentiation or specification to tuft-1 or tuft-2 subsets. This suggests that our *in vivo* observations are not driven by changes in tuft cell differentiation but may reflect changes in tuft cell activation state.

To exclude secondary involvement of immune cells and to pinpoint the intrinsic activation state of tuft cells under CD45 deletion, we compared the transcriptional states of WT and CD45-deficient organoids under non-treated (NT) and IL-13-stimulated conditions. Non-treated CD45-deficient organoids exhibited a transcriptional signature of 232 upregulated DEG that were shared with both IL-13-stimulated groups and absent from non-treated WT organoids. (**Fig. 5 E-F,** Venn diagram yellow and blue intersection). Since organoids include all epithelial lineage populations, the signature we obtained is not necessarily tuft cell-specific. To obtain a tuft-specific signature, we examined which of the 232 DE genes overlapped with upregulated DEG of tuft cells from CD45^ΔIEC^ compared with CD45^fl/fl^ mice (**Fig. 3 A**). We identified a shared tuft-associated signature of 40 genes (**Fig. 5 F, Table S4**). Representative genes within this signature included *Gata4, Chuk, Rac1, Pink1, Gclm*, *Selenoo*, and *Faf1*. Functional enrichment analysis broadly linked this gene set to response to stimulus (**Fig. S6 E**). More specifically, GO terms include negative regulation of intrinsic apoptotic pathway, protein dephosphorylation, and TNFα-NFκB signaling pathway (**Fig. 5 G**). Together, these data define a refined tuft cell-activation program that emerges either in the absence of CD45 at homeostasis or following IL-13 stimulation. These findings suggest that CD45 acts to maintain homeostasis by restraining an activation transcriptional program normally induced during IL-13-driven inflammation.

### CD45 deficiency is associated with increased STAT5 and reduced IL17RB abundance in tuft cells

We next sought to define candidate mechanisms by which CD45 can restrain tuft cell activation. CD45 is a protein tyrosine phosphatase that modulates cytokine receptor signaling, in part by dephosphorylating components of the JAK-STAT pathway and Src-family kinases (Irie-Sasaki et al., 2001; Saunders and Johnson, 2010). Given that tuft cells are chemosensory epithelial cells enriched for microbial, parasitic, taste, fatty-acid, and immune-sensing receptors (Feng et al., 2024), we examined candidate signaling pathways that could be regulated by tuft-specific CD45. Possible CD45 target candidates enriched in tuft cells at the RNA levels (Haber et al., 2017) included the direct dephosphorylation substrates of CD45, *Hck* and *Jak1* genes (Saunders and Johnson, 2010; Kumar et al., 2016). Tuft cells were also highly enriched for components of the IL-13 signaling pathway (*IL-13ra1*, *IL-4Ra*, and *Stat6*), the IL-25 receptor subunits *IL-17ra* and *IL-17rb*, and tuft effector regulators such as *Kit* and *Stat5a* (**Fig. 6 A**).

**Figure 6.**
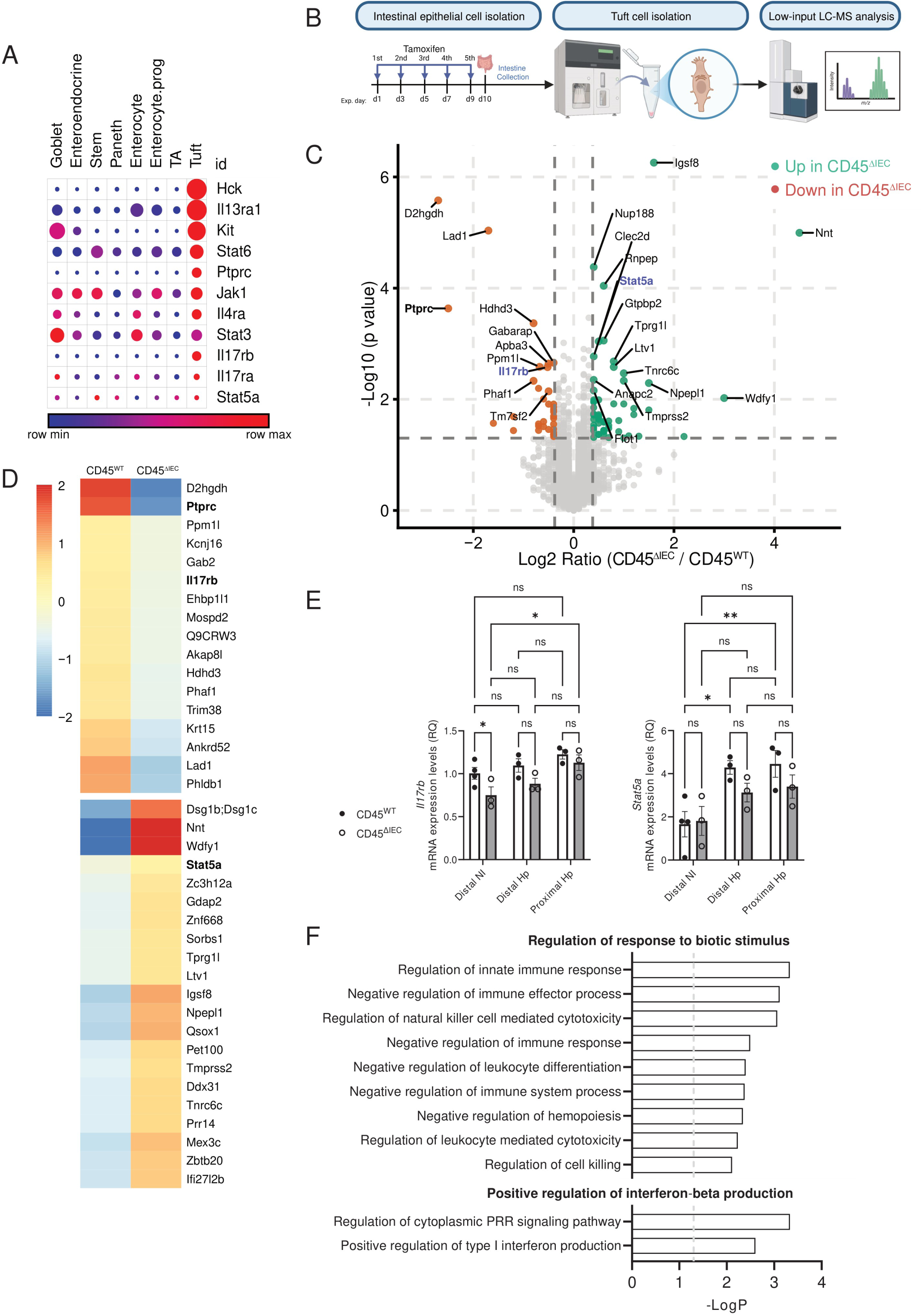
CD45 deficiency is associated with altered abundance of STAT5A and IL17RB in tuft cells. **(A)** Re-analysis of a published single cell RNA-sequencing dataset of the mouse small intestinal epithelium (Haber et al., 2017). Dot plot showing the expression of selected signaling and regulatory genes across intestinal epithelial cell populations. Dot size indicates the fraction of cells expressing each gene, and color indicates scaled gene expression. **(B)** Experimental design. CD45^ΔIEC^ and CD45^WT^ mice were treated with tamoxifen and distal SI was collected and processed, tuft cells were isolated via flow cytometry, and low-input liquid chromatography mass spectrometry was performed, n = 5-6 mice per group. Schematic created with BioRender. **(C)** Volcano plot of proteins identified by low-input liquid chromatography-mass spectrometry (LC-MS) in tuft cells isolated from CD45^ΔIEC^ and CD45^WT^ mice. Proteins increased in CD45^ΔIEC^ tuft cells are shown in green, and proteins decreased in CD45^ΔIEC^ tuft cells are shown in orange. Selected proteins are annotated. **(D)** Heatmap showing representative proteins with significantly altered abundance in CD45^ΔIEC^ and CD45^WT^ tuft cells identified by low-input LC-MS. **(E)** qPCR analysis of *Il17rb* and *Stat5a* expression in tuft cells isolated from the proximal and distal small intestine regions of non-infected (NI) or *Heligmosomoides polygyrus bakeri* (*Hp*)-infected CD45^ΔIEC^ and CD45^WT^ mice. Data are shown as mean ± SEM. Statistical significance was determined using two-way ANOVA followed by Tukey’s multiple-comparison test. \**P* ≤ 0.05; \*\**P* ≤ 0.01; ns, not significant. **(F)** Gene Ontology enrichment analysis of proteins with significantly increased abundance in CD45^ΔIEC^ tuft cells identified by low-input LC-MS.

Tuft cells are a rare epithelial cell type in the gut (±1-3%), making them challenging to analyze at the protein level, and even more so for unbiased post-translational modifications (PTMs). To overcome this, we isolated 500 tuft cells from CD45^ΔIEC^ or CD45^fl/fl^ mice and performed low-input liquid chromatography-mass spectrometry (LC-MS) to identify proteomic changes associated with CD45 loss (**Fig. 6 B-D**, **Methods**). The low-input quantitative proteomic analysis identified 5,776 proteins, including 33 downregulated and 55 upregulated (**Table S5**). Among the downregulated proteins, CD45 (Ptprc) showed reduced protein levels, further confirming the tuft-specific CD45 deletion. The downregulated group also included the IL-25 receptor subunit IL-17rb, which has been implicated as an attenuator of type 2 immunity. Moreover, tuft cell-specific loss of IL-17rb was previously reported to enhance type 2 responses (Feng et al., 2025) (**Fig. 6, C and D**). In addition, Stat5a abundance was increased in CD45-deficient tuft cells, and a recent study proposed that Stat5a phosphorylation can enhance type 2 immunity (Wang et al., 2025) (**Fig. 6, C-D)**

We next examined whether these proteomic changes were reflected at the transcript level by analyzing *Il17rb* and *Stat5a* mRNA expression in sorted tuft cells from CD45^ΔIEC^ and CD45^fl/fl^ mice (**Fig. 6 E**). Consistent with the protein-level changes, *Il17rb* expression was significantly reduced in CD45-deficient tuft cells at homeostasis (**Fig. 6 E**). By contrast, *Stat5a* expression was not altered by CD45 deletion at homeostasis, although *Stat5a* was significantly increased in tuft cells during *Hp* infection (**Fig. 6 E**). This discrepancy between STAT5A protein abundance and *Stat5a* mRNA expression may reflect post-transcriptional or post-translational regulation. Given that CD45 is a receptor-type protein tyrosine phosphatase, CD45 deletion in tuft cells may affect STAT5A abundance or signaling at the protein level rather than through changes in *Stat5a* transcription. Finally, pathway enrichment analysis of the upregulated proteins identified “regulation of response to biotic stimulus” and “positive regulation of interferon-beta production” as prominent categories, capturing multiple immune- and host-response-associated processes (**Fig. 6 F**). Overall, these results identify reduced IL-17rb and increased Stat5a abundance as CD45-associated signaling targets, which are implicated in promoting tuft effector activation. Whether these changes reflect direct CD45-dependent regulation or indirect downstream consequences of CD45 loss in tuft cells remains to be determined.

### IL-13 regulates CD45 expression in mouse tuft cells, with conserved expression in human tuft cells

After observing increased CD45 expression during *Hp* infection (**Fig. 4 B, Fig. S4 B**), we next asked which signals regulate its expression. In particular, it remains to be determined whether CD45 expression is induced directly by the parasite or by host-derived immune signals. IL-13, produced by ILC2s and Th2 cells, has previously been shown to act on Lgr5^+^ ISCs and promote differentiation toward tuft and goblet cell lineages (Gerbe et al., 2016; Howitt et al., 2016; von Moltke et al., 2016; Biton et al., 2018). Interestingly, our re-analysis of publicly available scRNA-seq data from mouse SI epithelium revealed a strong enrichment of the IL-13 response machinery in tuft cells, including IL13Ra, IL4Ra, and Stat6 (Haber et al., 2017), in line with the human organoid study of tuft cells (Huang et al., 2024) (**Fig. 7 A**). Furthermore, we observed a significant increase in *IL-13ra* expression in tuft cells isolated from *Hp*-infected mice (**Fig. 7 B**), suggesting that the observed CD45 increase may be mediated by tuft-intrinsic IL-13 signaling.

**Figure 7.**
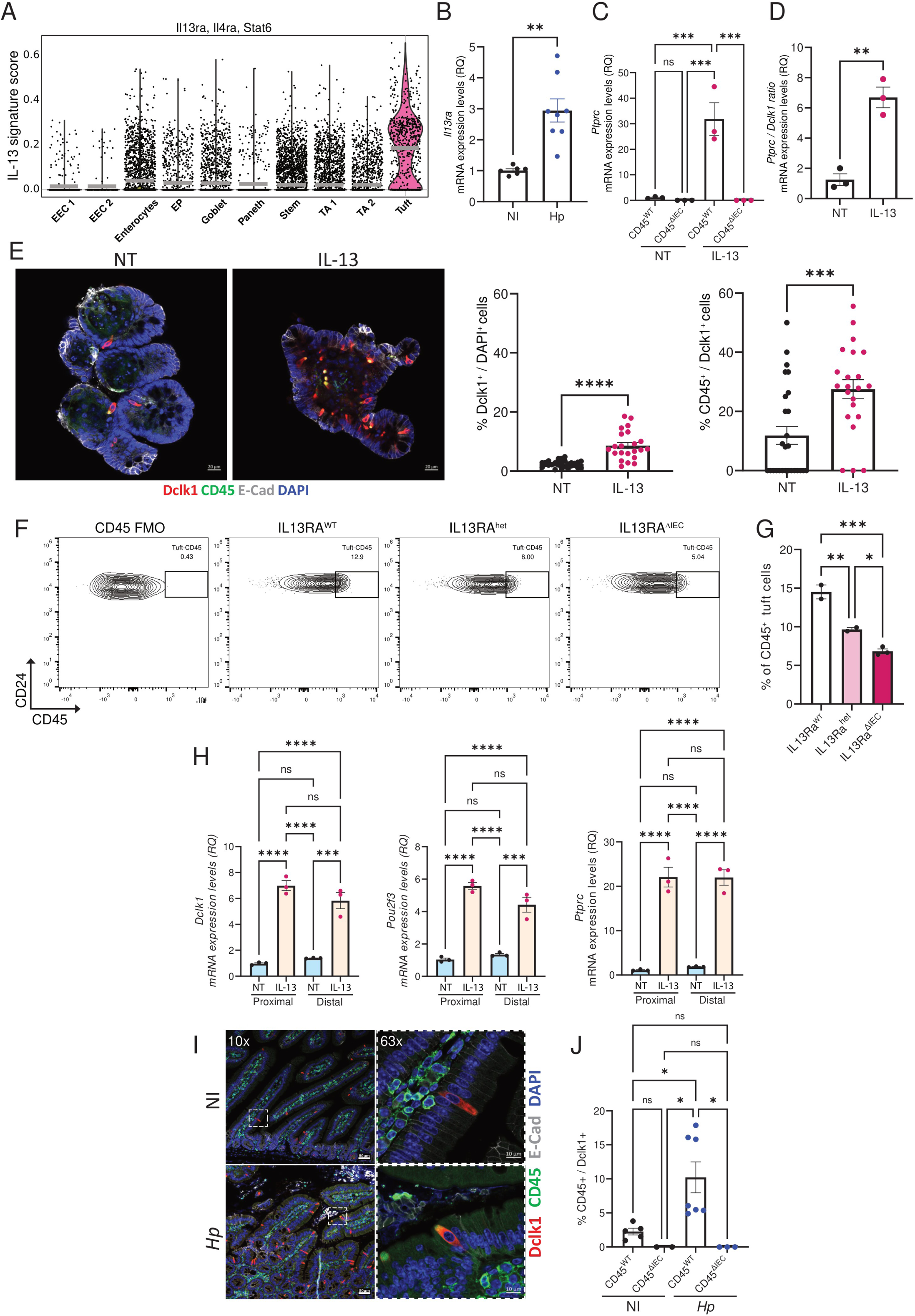
IL-13 induces CD45 expression in intestinal tuft cells. **(A)** IL-13 signaling signature score (*Il-13ra1*, *Il-4ra*, and *Stat6*) across intestinal epithelial cell populations from a published mouse small intestinal single-cell RNA sequencing dataset (Haber et al., 2017). **(B)** qPCR analysis of *Il-13ra1* expression in distal small intestinal tuft cells isolated from non-infected (NI) and *Heligmosomoides polygyrus bakeri* (*Hp*)–infected CD45^WT^ mice, n = 6-8 mice per group. **(C)** qPCR analysis of *Ptprc* expression in distal small intestinal organoids derived from CD45^ΔIEC^ and CD45^WT^ mice treated or non-treated (NT) with IL-13 for 72 h. n = 3 mice per group. **(D)** Relative *Ptprc/Dclk1* mRNA expression ratio in CD45^WT^ organoids cultured with or without IL-13, as measured by qPCR. **(E)** Left, representative immunofluorescence images of non-treated (NT) and IL-13-treated CD45^WT^ organoids stained for DCLK1 (red), CD45 (green), E-cadherin (gray), and DAPI (blue). Mid, quantification of DCLK1⁺ tuft cells presented as the percentage of DCLK1⁺ cells per DAPI⁺ cells; right, the frequency of CD45⁺ tuft cells expressed as the percentage of CD45⁺DCLK1⁺ cells per DCLK1⁺ cells. Each dot represents the average percentage per one organoid, n = 22-29 organoids per condition were quantified. **(F)** Representative flow cytometry analysis of CD45 expression in distal small intestinal tuft cells from *Il-13ra1*^WT^, *Il-13ra1*^het^ (fl/-), and *Il-13r*a^ΔIEC^ mice. **(G)** Quantification of CD45⁺ tuft cells in the genotypes shown in panel **F**. n = 2-3 mice per genotype. **(H)** qPCR analysis of *Dclk1*, *Pou2f3*, and *Ptprc* expression in matched proximal and distal small intestinal organoids cultured and treated or non-treated (NT) with IL-13 for 72 h. n = 3 organoid wells per condition. **(I)** Representative immunofluorescence images of proximal small intestine from non-infected (NI) and *Hp*-infected mice stained for DCLK1 (red), CD45 (green), E-cadherin (gray), and DAPI (blue), scale bar 50μm. Insets show high-magnification images of representative tuft cells, scale bar 10μm. **(J)** Quantification of CD45⁺ DCLK1⁺ out of DCLK1^+^ tuft cells in proximal small intestinal tissue from non-infected and *Hp*-infected mice. Quantification was assessed by 10 FOV per mouse, n = 2-7 mice per group. Data are shown as mean ± SEM. Statistical significance was determined using an unpaired two-tailed Student’s *t* test (**B**, **D**, **E**) or one-way ANOVA followed by Tukey’s multiple-comparison test (**C**, **G–J**). \**P* ≤ 0.05; \*\**P* ≤ 0.01; \*\*\**P* ≤ 0.001; \*\*\*\**P* ≤ 0.0001; ns, not significant.

We therefore asked whether IL-13, a major driver of tuft cell expansion during *Hp* infection, also regulates CD45 expression to initiate a negative-feedback loop of tuft cell activation. For this purpose, we used distal SI organoid cultures, which provide a controlled epithelial system in the absence of parasite-derived inputs. qPCR analysis showed an approximately 30-fold increase in *Ptprc* expression in IL-13-treated organoids compared with non-treated controls (**Fig. 7 C**, **Methods**). Because IL-13 promotes tuft cell differentiation, we also assessed canonical tuft cell markers. Whereas *Dclk1*, *Pou2f3,* and *IL-25* increased by approximately 5-6-fold following IL-13 treatment (**Fig. 5 B**), *Ptprc* increased to a greater extent, exceeding the induction observed for *Dclk1* by approximately 6.7-fold (**Fig. 7 D**). These data suggest that the increase in *Ptprc* expression cannot be attributed solely to IL-13-induced expansion of the overall tuft-cell population. Consistent with this, IF imaging of organoids showed that while IL-13 treatment significantly increased the proportion of DCLK1^+^ tuft cells from 2.4% to 8.6%, DCLK1^+^CD45^+^-expressing cells elevated from 11.9% to 27.5% within the tuft-cell population (**Fig. 7 E**).

To test the contribution of IL-13 signaling to CD45 expression in tuft cells *in vivo*, we utilized an epithelial-specific *IL13ra1* deletion mouse model (IL-13ra1^ΔIEC^) (**Methods**) and quantified the CD45^+^ tuft-cell frequency in the IL-13ra1^ΔIEC^ mice compared to corresponding controls (**Fig. 7 F-G)**. We observed a genotype-dependent reduction in CD45^+^ tuft-cell frequency, with a partial decrease from 14.5% to 9.6% in *IL-13Ra1* heterozygous mice and a larger decrease to 6.8% in homozygous *IL13ra1-*deleted mice. These findings are consistent with a gene-dosage effect of epithelial IL-13 signaling on CD45 expression. Notably, overall tuft-cell frequency was unchanged in IL13RA1^ΔIEC^ mice (**Fig. S7 A**).

IL-13 induced *Ptprc* expression in distal SI and organoids; we next asked whether this effect also takes place in the proximal SI. To address this, we established organoids from matched proximal and distal SI crypts from the same mice and stimulated them with IL-13 for 72h (**Fig. 7 H**). *Dclk1* and *Pou2f3* increased by approximately 5-6-fold in both compartments. *Ptprc* expression likewise increased by approximately 20-fold in both proximal and distal IL-13-treated organoids (**Fig. 7 H**). These results suggest that CD45 expression is induced by IL-13 without regional preference. We then examined proximal SI tissue during *Hp* infection to determine whether type 2 immune activation is associated with CD45 expression *in vivo*. In this setting, the proportion of CD45^+^ tuft cells was significantly higher in infected mice than in controls, increasing from 2.2% to 10.2% (**Fig. 7 I-J**). Together, these data support IL-13 as a regulator of tuft-cell CD45 expression across organoid and *in vivo* models, in both the proximal and distal SI. This IL-13-mediated CD45 expression prevents excessive activation of type 2 immunity in the intestine in an IL-13-dependent negative feedback loop.

To determine whether tuft cell CD45 expression is conserved across species, we examined publicly available scRNA-seq datasets from healthy adult human intestines (Hickey et al., 2023). Similarly to mice, the ileal epithelium of adult humans showed the highest enrichment for the IL-13 sensing signature in tuft cells (IL-*13RA1*, *IL13RA2, IL4R,* and *STAT6*, **Fig. S7 B**). In addition, *PTPRC* expression showed enrichment in tuft cells, consistent with the pattern observed in mice (**Fig. S7 C**). Finally, we examined the human SI regional distribution of tuft *PTPRC* expression. In line with our observations in mice, *PTPRC* expression was enriched in ileal tuft cells (**Fig. S7 D**) (Hickey et al., 2023). Thus, human tuft cell data suggest that a similar tuft cell-specific CD45 regulatory axis operates in humans, although this requires further validation.

## Discussion

A growing body of evidence points to region-specific regulation of type 2 immunity along the small intestine. In the proximal SI, IL-25 alone is insufficient to initiate ILC2 activation and downstream type 2 immunity in response to *Hp*. Instead, tuft cell-derived leukotrienes are required for this response, whereas in the distal SI, leukotrienes are dispensable (McGinty et al., 2020). Despite this distinction, type 2 immunity in the distal SI remains tightly controlled. Moreover, although the distal SI is rich in succinate, a microbiota metabolite that can activate the tuft cell-mediated type 2 immunity circuit (Nadjsombati et al., 2018; Schneider et al., 2018), excessive type 2 immune activation remains in check. These observations suggest that other mechanisms may actively restrain type 2 immunity in the distal SI under homeostasis. Given CD45’s known role as a threshold modulator of immune responses in immune cells, we hypothesized that tuft cell CD45 expression may restrain tuft cell responses at steady states.

In this study, we identify epithelial CD45 as a tuft cell-specific regulator that restrains type 2 immunity in the distal small intestine both at steady state and under helminth infection. In the absence of epithelial CD45, we observed a tuft cell-activated mode that led to an overall epithelial inflammatory landscape, including extensive transcriptional changes in IECs and expansion of type 2-associated immune populations, with accumulation of eosinophils at homeostasis and of ILCs and Th2 cells during infection.

Within the established type 2 circuit, IL-13 drives tuft and goblet cell expansion, thereby propagating the tuft-ILC2 positive feedback loop (**Fig. S8**). Our data suggest that IL-13 also induces CD45 expression in tuft cells, which in turn attenuates further amplification of type 2 immunity (**Fig. S8**). Consistent with this model, CD45 expression increased in both IL-13-treated organoids and *Hp*-infected mice. Moreover, epithelial *IL-13ra1* expression was required for CD45 induction *in vivo*, and the magnitude of CD45 expression depended on the number of functional IL-13ra alleles. Further supporting this IL-13-dependent regulatory axis, we found that *IL-13ra1* expression is also increased in *Hp*-infected tuft cells. By contrast to *Hp* infection, where tuft cell expansion is dependent on IL-13 expression, epithelial deletion of IL-13ra1 did not alter steady-state tuft cell abundance, suggesting that epithelial IL-13 sensing is required for tuft cell activation and CD45 induction, rather than for baseline tuft cell differentiation under homeostasis. Likewise, CD45 deficiency did not alter overall tuft cell differentiation or the balance between tuft-1 and tuft-2 states, supporting a role for CD45 in tuning tuft cell activation rather than lineage specification. In line with these findings, we propose a model in which during *Hp* infection, IL-13 initially promotes propagation of the type 2 immune circuit by driving tuft cell differentiation, but as IL-13 levels rise in the intestine, IL-13ra levels also increase in tuft cells, inducing the tuft-intrinsic type 2 negative regulator CD45, and thereby restraining further amplification of the response (**Fig. S8**).

While our work centers on tuft cell regulation in the distal SI, we also asked whether CD45 restrains tuft responses in the proximal SI. We found that both IL-13 stimulation and *Hp* infection induced CD45 expression in proximal SI tuft cells, indicating that this regulatory pathway is not restricted to the distal intestine. The regional difference observed at baseline may reflect differences in tonic exposure to activating cues. Distal SI tuft cells are exposed to higher levels of luminal and microbiota-derived metabolites, including succinate, which may activate them. Therefore, inhibitory mechanisms, such as CD45 expression in a subset of tuft cells, are required to prevent hyperactivation of the Th2 response in the distal region. This molecular switch may explain why approximately 20% of distal tuft cells express CD45 at homeostasis, whereas proximal tuft cells show little to no detectable expression under baseline conditions. During infection, however, proximal tuft cell CD45 expression rises to a similar level, indicating that regional CD45 expression is shaped by the inflammatory context rather than by a fixed intestinal identity. This model provides a plausible explanation for the regional distribution of tuft cell CD45 expression observed in our study.

Several recent studies have identified epithelial feedback mechanisms that restrain type 2 immunity by limiting IL-13-driven epithelial remodeling. Oyesola et al. showed that tuft-cell-derived PGD2 can act through CRTH2 on IECs to counteract type 2 cytokine-driven epithelial reprogramming, including *IL-13ra1* expression and secretory lineage differentiation (Oyesola et al., 2021). Similarly, Lindholm et al. identified an IL-13-induced BMP feedback loop that limits tuft cell hyperplasia by restricting Sox4-associated tuft progenitor differentiation (Lindholm et al., 2022). Thus, in these models, suppression of type 2 immunity is closely linked to altered epithelial differentiation, secretory lineage differentiation, or tuft cell expansion. In contrast, our data suggest that CD45 primarily regulates tuft-cell effector pathways rather than tuft cell differentiation. One tuft intrinsic effector pathway that regulates type 2 immunity is Stat5a, which Wang et al. implicated in tuft cell TSLP production and Th2 activation (Wang et al., 2025). In line with this functional connection, we observed higher levels of Stat5a protein in tuft cells lacking CD45, indicating that CD45 may play a role in restraining Stat5a protein expression. Additionally, we observed increased *Stat5* expression in tuft cells following *Hp* infection, suggesting that Stat5a may be involved in tuft cell activation during type 2 inflammation. These findings imply that the elevated abundance of Stat5a during infection and in epithelial-specific CD45-deficient conditions could contribute to a heightened type 2 immune response. We also observed reduced protein expression of the IL-25 receptor subunit IL-17rb, which has been shown to limit excessive IL-25-driven ILC2 stimulation (Feng et al., 2025), suggesting altered regulation of the tuft cell-ILC2 circuit in the absence of CD45 in tuft cells. Together, our findings suggest that CD45 may tune multiple tuft cell effector pathways rather than acting through a single downstream node. These findings position Stat5a and IL-17rb as candidate components of a CD45-mediated tuft cell program that attenuates type 2 immune output (**Fig. S8**).

Using the CD45 depletion model, we identified a tuft activation signature that is negatively regulated by the CD45-mediated pathway. This tuft cell activation signature is enriched for defense responses to infection and the NFκB inflammatory pathways. The activated tuft state observed in CD45-deficient tuft cells has a significant functional role during *Hp* infection, contributing to enhanced parasite expulsion, as reflected by the lower worm burden in these mice.

Overall, our study provides a comprehensive characterization of a non-hematopoietic CD45-dependent tuft cell-intrinsic pathway that restrains type 2 immunity. These findings are consistent with the concept of disease tolerance, described by Medzhitov et al. as a host strategy that limits the adverse effects of the host immune response during infection while maintaining effective immune protection (Medzhitov et al., 2012). In line with this idea, our findings suggest that tuft cell CD45 limits excessive type 2 immune activation while preserving effective control of *Hp* infection. Finally, our findings support a new model in which IL-13 both amplifies and attenuates type 2 immunity via the chemosensory tuft cells, thereby acting as an immune rheostat.

## Acknowledgments

We thank Prof. Steffen Jung, Prof. Rony Seger and Prof. Ronen Alon from the Weizmann Institute of Science, and Prof. Yehudit Bergman from the Hebrew University for insightful discussions. We thank Dr. Shira Albeck for cloning the in house Tn5. We thank Polina Gardchenko, Eva Varshavski, Hadar Klimovski, Maayan Donitza and Einav Haya Hirik for genotyping mice. We thank Muriel Chemla for assisting with sequencing. M.B. holds the Ernst and Kaethe Ascher Career Development Chair. This study was supported by research grants from the Center for New Scientists at the Weizmann Institute of Science, the Israel Science Foundation (grant No. 1587/20 and 3775/25), the Helen and Martin Kimmel Institute for Stem Cell Research at The Weizmann Institute of Science, and the Minerva Foundation, with funding from the Federal German Ministry for Education and Research, the Moross Integrated Cancer Center, the Israel Ministry of Science (IMOS, Grant No. 4631), the Dr. Gilbert S. Omenn and Martha A. Darling Weizmann Institute - Schneider Hospital Fund for Clinical Breakthroughs through Scientific Collaborations, a research grant from the Another Light Foundation, the Abisch-Frenkel RNA Therapeutics Center, a research grant from the Shimon and Golde Picker, and a research grant from the Herbert K. Bennett Charitable Fund, Dwek Institute for Cancer Therapy Research and the Belle S. and Irving E. Meller Center for the Biology of Aging.

## Author contributions

C.S. and M.B. conceived the study, designed experiments, and interpreted the results; C.S. carried out all experiments with the help of S.L., A.H.M., V.H. and V.R.; C.S. performed RNA-seq experiments; C.S. and A.S.P. designed and performed the computational analysis with the assistance of B.T.; C.S., J.L and Z.S performed MLN B cell analysis; C.S. and S.L. performed *Hp* infections; C.S performed organoid stimulation experiments with the assistance of S.L.; Y.L. performed proteomics profiling; E.V. assisted with immunofluorescence imaging and analysis; N.R. assisted with preliminary experiments; N.T. and D.K.A. assisted with IL13Ra^ΔIEC^ mice experiments, C.S., I.O. and S.B.D. designed CRISPR Cas-9 guides to generate CD45^fl/fl^ mice; R.H.K. generated transgenic CD45^fl/fl^ mice; R.K.G. and D.K.A. provided *Hp* larvae and parasite culture guidance. A.M. provided IL13Ra KO mice, M.B. supervised this study; C.S. and M.B. wrote the manuscript, with input from all authors.

## Materials & Methods

### Mice

All mouse work was performed in accordance with the Institutional Animal Care and Use Committees (IACUC) of the Weizmann Institute of Science (IACUC numbers 07731023-2, 03080323-2). Wild type (WT) (C57BL/6J; 000664), Villin-CreER^T2^ (020282), and Chat GFP (007902) mice were purchased from the Jackson Laboratory. Both male and female age-matched mice, ranging from 8 to 14 weeks of age, were used for all experiments in this study. Littermates of the same genotype, sex, and age were randomly assigned to experimental groups. All mice were housed under specific-pathogen-free (SPF) conditions at the Weizmann Institute animal facilities. Villin-CreER^T2^ x IL-13ra (IL-13ra^ΔIEC^) mice were kindly gifted by Dr. Danielle Karo-Atar from Ben-Gurion University of the Negev.

CD45 floxed mice (background C57BL/6JOlaHsd from Harlan) were generated ‘in house’ using CRISPR. In brief, CRISPR guides for Cas9 were chosen using several CRISPR design tools including: the MIT CRISPR design tool (Hsu et al., 2013) and sgRNADesigner (Doench et al., 2016), in the Benchling implementations (www.benchling.com) and CRISPOR (Concordet and Haeussler, 2018). Flox sites were designed to flank the promoter, exon 1 and exon 2 of the PTPRC gene, which should eliminate all gene isoforms when crossed with Cre lines. Then, transgenic mice were generated by Dr. Rebecca Haffner-Krausz (transgenic mice unit, Weizmann institute). Synthetic crRNA, tracrRNA, HDR oligos and Cas9 nuclease were purchased from IDT. CD45 flox mice were backcrossed with C57BL/6 WT mice for at least 3 generations to avoid off-target effects. Finally, mice with both flox sites on the same allele were bred with Vil-CreER^T2^ mice to obtain homozygous CD45 flox mice with Cre under the Villin-CreER^T2^ promoter (CD45^ΔIEC^). Guides against the *Ptprc* gene:

Guide 1: GGTTATTTCCATGTACAGAA

Repair oligo guide 1:

TTTGGGCAAATATGAAATAATATGATTCCTTGAGGCTTTAGAATTTTATTACCTGGAATTTAAGTTGTAAGGTTATTTCCATGTACACTCGAGATAACTTCGTATAGCATACATTATACGAAGTTATGAAAGGGGGATGTGCAAGTTTGACCCTGCAGCTCCTTGAAATCAAAGTAAGAGGAAAACAAAGTTTTAAGCAT

Guide 2: AATCGGAAAATTGCCCCGAG

Repair oligo guide 2:

AGAATTATAGAGAAGAATTGAAGATTGTACATTTTATCATATATTGGCATATTTCCACTGTAAGTCCCCACTCCTCGAGATAACTTCGTATAGCATACATTATACGAAGTTATGGGGCAATTTTCCGATTTTCTTTAAATGGGTGAGAAAGAACATTAAGGTACCTTCAGATTGTAGAGCAAGTCACCCCCAAATTATAA

All genotyping primers are available in **Supplementary Table S6**.

### Conditional deletion of CD45 in tuft cells

To initiate CD45 deletion, mice were administered 5 intraperitoneal injections of Tamoxifen every other day (2 mg per 20 g body weight) to induce Cre-mediated excision of the floxed region of *Ptprc* to confer its deletion. Mice were sacrificed 10 days after the first injection unless specifically noted otherwise.

#### Mice *Heligmosomoides polygyrus Bakeri* infection

*Heligmosomoides polygyrus bakeri* was propagated in-house as previously described (Johnston et al., 2015). Briefly, feces from infected donor mice were collected starting 4 weeks post-infection and seeded onto sterile water-moistened Whatman paper in 15-cm plates. Plates were maintained under moist conditions, and third-stage larvae (L3) were collected 10–14 days after seeding by washing the Whatman paper with sterile water. L3 were pelleted by centrifugation at 300 × g for 3 min, resuspended in sterile water, and counted by microscopy. L3 were kept in 5ml sterile water at 4°C for up to 6 months. Larva was counted and motility was confirmed before infection. *Hp* larvae were kindly gifted by Prof. Richard Grencis from Manchester University and Dr. D. Karo-Atar from Ben-Gurion University of the Negev.

Experimental mice were infected with 200 motile L3 larvae in 100 µL sterile water by oral gavage. Infected mice were maintained under specific-pathogen-free conditions at the Weizmann Institute of Science in accordance with IACUC protocols 03080323-2 and 07731023-2. Mice were euthanized 10 or 14 days post-infection.

### Epithelial cell dissociation and crypt isolation

For all mice (unless noted specifically otherwise), crypts were isolated from the distal part of the SI. The SI was extracted and rinsed in cold Phosphate buffered saline (PBS). The tissue was opened longitudinally and sliced into small fragments roughly 2 mm long followed by incubation with 20mM EDTA-PBS on ice for 90 min. Then, the tissue was shaken vigorously, and the supernatant was collected as fraction 1 in a new conical tube. The tissue was incubated in fresh PBS, and a new fraction was collected every 5 min. Fractions were collected until the supernatant consisted almost entirely of crypts. The fractions 3-4 (enriched for crypts) were filtered through a 70 µm filter, centrifuged at 400g for 5 min, and dissociated with TrypLE Express (Gibco) for 1 min and 30 seconds at 37°C. The single-cell suspension was then passed through a 40μm filter and stained with fluorescence-activated cell sorting (FACS) antibodies and sorted with SH800 Sony sorter for subsequent analysis. FACS sorting was performed for qPCR, bulk RNA-seq, and proteomics (see below).

### Lamina Propria immune cell isolation

After villus and crypt isolation, immune cells from the Lamina Propria (LP) were isolated enzymatically by incubating the small intestine with Liberase TM (100 μg/mL, Sigma) and DNase I (10 μg/mL, Sigma) for 45 min at 37°C while rotating. The tissue was then meshed through the 70 μm filter, and cells were centrifuged at 400g for 5 min. Immune cells were stained for either flow cytometry analysis or FACS.

### Mesenteric lymph nodes isolation

Mesenteric lymph nodes (MLN) were processed on ice in PBS x 1. MLNs were then passed through 70 µm strainers. Cells were washed (400 × g, 7 min), resuspended in PBS x1 and stained with an antibody cocktail in PBS x1 (described below) for 30 minutes on ice. Then, cells were washed with PBS x1, centrifuged at 400 × g, 7 min, resuspended in FACS buffer (PBS containing 2% fetal bovine serum and 1 mM EDTA) and kept on ice in the dark until flow cytometry analysis.

### Flow cytometry analysis of epithelium and lamina propria cells

Single-cell suspensions were prepared as described above. Epithelial cells were stained with antibodies against EpCAM (BioLegend, G8.8), CD45 (BioLegend, 30-F11), CD24 (BioLegend, M1/69), SiglecF (BioLegend, QA20A11) and Dapi-NucBlue (ThermoFisher, cat no. R37605) for 30’ on ice. The CD45^+^ tuft gate was determined using a CD45 fluorescence minus one (FMO) sample.

LP immune cells were stained with Zombie Aqua Fixable Viability dye (BioLegend cat 423102) and antibodies against CD45 (BioLegend, 30-F11), IL-17rb (BioLegend, 9B10), CD11b (BD, M1/70), SiglecF (BioLegend, QA20A11), CD3 (BioLegend, 500A2), CD4 (BD, GK1.5), CD8 (BD, 53-6.7), TCRβ (BioLegend, H57-597), Lineage negative(Nevo et al., 2024) on APC-cy7 [(CD127 (BioLegend, A7R34), CD19 (BioLegend, 6D5), NK1.1 (BioLegend, PK136), GR-1 (BioLegend, RB6-8C5), TCRγ/δ (BioLegend, GL3), CD11c (BioLegend, N418)], GATA3 (BioLegend, 16E10A23), FOXP3 (BD, R16-715), RORɣt (BD, Q31-378), T-bet (BioLegend, 4B10). Intracellular staining was performed using the True-Nuclear™ Transcription Factor kit (BioLegend, cat. No. 424401) according to the manufacturer’s guidelines. Analysis was done on a CytoFLEX S flow cytometer or a Sony SH800 sorter. Further analysis was performed using FlowJo v10.10.1.

### Flow cytometry analysis of mesenteric lymph nodes immune cells

For the B cell FACS panel, cells were stained with antibodies against CD45 (BioLegend, 30-F11), B220 (BioLegend, clone RA3-6B2), CD38 (Invitrogen, clone 90), FAS/CD95 (BD Biosciences, clone Jo2), GL-7 (BioLegend, clone GL7), CD138 (BioLegend, clone 281-2), IgA (BioLegend, clone RMA-1), IgM (BioLegend, clone MA-69), IgG1 (BioLegend, clone RMG1-1), and IgG2a/b (Miltenyi Biotec, clone X-57).

For the T cell FACS panel, cells were stained with antibodies against CD45 (BioLegend, 30-F11), CD4 (BD, GK1.5), CD8 (BD, 53-6.7), CD44/Pgp-1/Ly-24 (BD Biosciences, clone IM7), CD62L (BioLegend, clone MEL-14), CXCR5 (BioLegend, clone L138D7), PD1 (BioLegend, clone 29F.1A12) and B220 (BioLegend, clone RA3-6B2).

Analysis was done on CytoFLEX S flow cytometer. Further analysis was performed using FlowJo v10.10.1.

### Cell sorting

For the bulk population, FACS (SH800 Sony) was used to sort 1,000-50,000 cells into an Eppendorf tube containing 50 μL TCL buffer (QIAGEN) solution with 1% β-mercaptoethanol (Sigma-Aldrich). To enrich Tuft cells, cells were gated on DAPI^neg^ EpCAM^+^ CD45^neg-mid^ CD24^hi^ FSC-A^low^ in all results except for **Figure 1**, for which cells were gated on DAPI^neg^ EpCAM+ CD45^neg-mid^ CD24^hi^ SiglecF^+^. For bulk RNA sequencing, the tubes were centrifuged, immediately flash-frozen on dry ice, and stored at -80°C until ready for RNA isolation and library preparation.

For low-input liquid chromatography mass spectrometry (LC/MS) analysis, samples were flash-frozen in LC/MS lysis buffer (0.2% n-dodecyl β-D-maltoside (DDM), NaCl 150mM) and kept at -80°C until ready for protein isolation.

### Smart-seq2 bulk RNA-seq libraries

Libraries were prepared using a modified SMART-Seq2 protocol (Picelli et al., 2013, 2014b). In brief, RNA lysate clean-up was performed using RNAClean XP beads (Agencourt), followed by reverse transcription with Maxima Reverse Transcriptase (Life Technologies) and whole-transcription amplification with KAPA HotStart HIFI 2 × ReadyMix (Kapa Biosystems) for 18 cycles. whole-transcription amplification products were purified with Ampure XP beads (Beckman Coulter), quantified with Qubit dsDNA HS Assay Kit (ThermoFisher) and assessed with a high-sensitivity DNA chip (Agilent). RNA-seq libraries were constructed from purified whole-transcription amplification products using Nextera XT DNA Library Preparation Kit (Illumina, FC-131), or using our in-house Tn5 transposase (Picelli et al., 2014a). The libraries were sequenced on an Illumina NovaSeq 6000 for sequencing data presented in Fig. 3, and NovaSeq X on the rest of the sequencing data.

### Tn5 cloning and expression

Cloning and production of in-house Tn5 (iTn5) was performed by Dr. Shira Albeck from the Department of Life Sciences Core Facilities according to previously published protocols (Picelli et al., 2014a; Hennig et al., 2018).

### Quantitative PCR

For quantitative PCR (qPCR), cDNA from bulk population libraries was diluted to a final concentration of 0.5 ng/μL. Fast SYBR green master mix (Thermo-Fisher Scientific) was used to perform qRT–PCR per the manufacturer’s instructions. Data analysis was performed with QuantStudio 12K flex software (Thermo-Fisher Scientific) based on the ΔΔCT method. All target genes were standardized to the endogenous reference gene Ubiquitin C (*Ubc*). For the primer sequences used, see **Supplementary Table S7.**

### Low-Input Liquid Chromatography Mass Spectrometry

#### Sample collection for low-input mass-spectrometry

Cells were FACS sorted directly into 0.2% n-dodecyl β-D-maltoside (DDM) 150mM NaCl (final concentration). The lysates were reduced with 6.25 mM dithiothreitol for 30 min followed by alkylation with 12.5 mM iodoacetamide in the dark for additional 30 min. Samples were then digested with 30ng of trypsin for 4 h at 37 °C in a PCR thermocycler. The digested peptides were acidified with 0.1% TFA, then stored in -20°C until analysis.

#### Liquid chromatography

Each sample was loaded using nano-Ultra Performance Liquid Chromatography (nanoElute2, Bruker, Germany). The mobile phase was: A) H2O + 0.1% formic acid and B) acetonitrile + 0.1% formic acid. The peptides were separated using an Aurora column (75μm ID x 25cm, IonOpticks) at 0.3 µL/min. Peptides were eluted from the column into the mass spectrometer using the following gradient: 2% to 35%B in 60 min, 35% to 95%B in 0.5 min, maintained at 95% for 4.65 min.

#### Mass Spectrometry

The nanoUPLC was coupled online to a quadrupole time-of-flight mass spectrometer (timsTOF Ultra, Bruker) using the CaptiveSpray nanoESI source (Bruker Daltonics, Germany). Data was acquired in Parallel Accumulation–Serial Fragmentation combined with Data-Independent Acquisition (DIA-PASEF) mode (Meier et al., 2020). MS1 mass range was set to 100-1,700 Th. MS1 Ion mobility range was set to 0.70-1.30 1/K0, ramp time of 166 msec. 27 windows of 26 Da with 1 Da overlap were set over a range of 348-1024 Th, cycle time of 1.2 sec.

#### Data processing

Raw data was processed with DIA-NN version 2.2 (Demichev et al., 2020). The data was searched using a predicted library based on the SwissProt mouse proteome database (January 2025 version) in addition to a common contaminants list. Carbamidomethylation of C was set as a fixed modification. Oxidation of M, and protein N-term acetylation were set as variable modifications (maximum 2). Up to two mis-cleavages were allowed, proteotypicity was set to ‘protein names from FASTA’. Precursor false discovery rate was set to 1%. The protein intensities were used for further calculations using Perseus v1.6.2.3. The protein intensities were log2-transformed and only proteins that had at least 70% valid values in at least one experimental group were kept. The remaining missing values were imputed by a random low range distribution. Student’s t-tests were performed between the relevant groups to identify significant changes in protein levels.

### Histochemistry

Tissues were fixed for 16-24 hours in formalin, embedded in paraffin, and cut into 5μm-thick sections. Sections were deparaffinized with standard techniques. For H&E staining, slides were stained with Hematoxylin for 1 minute, washed, and then stained with Eosin for 45 seconds. For PAS and Alcian blue staining, tissue was stained with Alcian blue and periodic acid-Schiff reagents kit (ScyTek Laboratories, cat no. APS-1) on deparaffinized slides with standard techniques, using 3% acetic acid for 3 minutes, then Alcian blue solution (Alcian blue, pH 2.5) for 30 min, rinsed in tap water for 5 min, and oxidized in periodic acid (0.5%) for 10 min, followed by rinsed in running tap water for 5 min, and stained in Schiff reagent as a counterstain (cancer diagnostics) for 10 min. Nuclei were stained with PureView™ Mayers Hematoxylin for 50 seconds. All slides were washed, dehydrated and mounted with Sub-x mounting medium (Leica).

### Immunofluorescence (IF)

Tissues were fixed for 14 hours in formalin, embedded in paraffin, and cut into 5-μm-thick sections. Sections were deparaffinized with standard techniques and incubated with primary antibodies overnight at 4°C, followed by incubation with secondary antibodies at room temperature for 45 min, and Hoechst 33342 (1:1000) (TargetMol, T5840). Slides were mounted with Fluoromount-G (SouthernBiotech) and sealed. Primary antibodies used for immunofluorescence include mouse anti-E-cadherin (1:100, BD Biosciences Clone 36), rat anti-Ki67 (1:100, Invitrogen SolA15), Rabbit anti-DCLK1 (1:300, Abcam ab31704), Rat anti-CD45 (1:200, 30-F11, Biolegend 103101). Secondary antibodies include Alexa Fluor 488-, 568-and 647-conjugated (1:400, Abcam).

### Image analysis

Images of tissue sections and organoids were taken with a confocal microscope LSM900 confocal microscope (Zeiss). Scale bars were added to each image using the confocal Zen analysis software (Zeiss). Images were overlaid and visualized using Zen analysis software (Zeiss). Organoids were analyzed with FIJI software.

### Image quantification

Quantification of IFA images from all tissues was performed by staining for E-Cadherin (BD Biosciences) to mark cell borders and for DAPI to visualize nuclei. Cells were manually counted based on immunofluorescence staining specific to each cell type. Tuft cells were identified by DCLK1^+^ staining and were quantified along the crypt-villus axis. CD45^+^ Tuft cells were identified by co-staining of CD45^+^ with DCLK1^+^, CD45^+^ DCLK1^-^ cells were restricted to the LP and were not counted. Proliferative cells (Ki67^+^) were counted along the villus-crypt axis. For each quantification, at least 10 intact, longitudinally oriented crypts per tissue section were analyzed. Quantification of histochemical images of goblet cells was performed using AB-PAS staining. For each quantification, at least ten randomly intact and longitudinally oriented crypts-villi per tissue section were analyzed.

### Crypt seeding for spheroid establishment and expansion

Following crypt isolation from different mouse models (WT, CD45^ΔIEC^, IL-13ra^ΔIEC^), the number of crypts was estimated using a bright field optical microscopy (EVOS™ M5000 Imaging System). About 500 crypts were embedded in 20 μL Matrigel™ (Corning) with 1µM Jagged-1 peptide (Ana-Spec). After 20 min at 37 °C in a humidified atmosphere at 5% CO_2_, 500ul of L-WRN conditioned media (C.M.) supplemented with 10 µM Y-27632 Dihydrochloride (Biogems, cat. no 1293823)10 µM Y27632 were added and incubated at 37 °C with 5% CO_2_.

### Organoid differentiation, culturing, and collection for downstream analysis

Once sufficiently expanded, spheroids were passaged and Matrigel domes were cultured in complete organoid medium based on Advanced DMEM/F12 supplemented with glutamine and penicillin–streptomycin. Complete organoid medium contained EGF (50 ng/mL; PeproTech, AF-100-15), Noggin (200 ng/mL; PeproTech, 250-38), R-spondin 1 (750 ng/mL; Sino Biological, 50316-M08H), Y-27632 (10 µM; Sigma-Aldrich, Y0503), N-acetylcysteine (NAC; 1 µM; Sigma), N2 supplement (1×; Invitrogen, 17502048), and B27 supplement (1×; Invitrogen, 0080085SA). For routine passaging, culture medium was removed, and organoids were collected from Matrigel domes by scraping in cold PBS. Organoids from each condition were pooled and kept on ice. Residual Matrigel was removed as much as possible without excessive sample loss, and organoids were mechanically dissociated by pipetting with cold PBS x1. Following centrifugation, the pellet was resuspended in Matrigel and plated as 10-20-µl domes. Plates were inverted during Matrigel polymerization, and complete organoid medium was added after 30 min. For the collection of organoids for bulk RNA-seq or qPCR analysis, organoids were washed extensively with PBS x1 to remove Matrigel and lysed directly in 100-200 μL TCL buffer (QIAGEN) solution with 1% β-mercaptoethanol (Sigma-Aldrich) before freezing at −80°C.

### Organoid IL-13 stimulation

Organoids were treated with IL-13 at 20ng/ml in the culture medium for 72h (Biton et al., 2018).

### Statistical analysis

Statistical analysis was performed with the Prism software (GraphPad software), R v4.4.1, R v4.3.1. The specific tests applied are detailed in the corresponding figure legends and computational methods.

### Computational analysis

#### Data and code Availability

The proteomics dataset is available through the ProteomeXchange Consortium via the PRIDE repository under accession number XXXXXX. Bulk RNA-seq datasets are available at the Gene Expression Omnibus at accession number **GSE339490**.

### Bulk RNA-sequencing

#### Read QC, trimming, and alignment/quantification

Samples were demultiplexed using bcl2fastq v2.20. Adapter and quality trimming were performed with cutadapt v2.7 (minimum length 25 nt, quality cutoff Q10). Read quality was assessed with FastQC v0.11.8. Trimmed reads were aligned to the Mus musculus reference genome GRCm39 with STAR v2.7.3a (with EndToEnd option) against the Ensembl release 110 gene annotation (GTF). Gene-level counts were obtained with HTSeq-count v0.12.4. Only uniquely mapped reads were used to determine the number of reads that map to each gene (intersection-strict mode).

#### Normalization and differential expression

Genes with fewer than 5 counts in all the samples were filtered. Differential expression was performed in R v4.2.2 using DESeq2 v1.38.3 with the betaPrior, cooksCutoff, and independentFiltering parameters set to False. Raw P values were adjusted for multiple testing using the procedure of Benjamini and Hochberg. Differentially expressed genes were determined by a p-adj of <0.05 and absolute fold changes >2 and a count of at least 30 in at least one sample. Heatmap plotting was performed with the ComplexHeatmap R package v2.14, using scaled, log2-transformed DESeq2-normalized counts for each gene. For gene-signature scoring, specific gene sets were curated and used. For each data set individually, raw DESeq2 counts were ranked per sample using rankGenes, and the singscore R package v1.18.0 was applied using the simpleScore function with default random background sampling (equal size). The resulting per-sample signature score was used for downstream visualization.

#### Gene Ontology enrichment analysis

Gene Ontology (GO) enrichment analysis was performed using Metascape (Zhou et al., 2019) to identify functional and pathway enrichment among differentially expressed genes or proteins. The analyses presented in Fig. 6 were performed in January 2026, those presented in Fig. 3-4 were performed in March 2026, and those presented in Fig. 5 were performed in May 2026.

For the RNA-sequencing datasets, differentially expressed genes were defined by an adjusted *P* value ≤ 0.05 and an absolute log2 fold change ≥ 1. Upregulated and downregulated genes were analyzed separately. When either gene set contained more than 200 differentially expressed genes, the 200 statistically significant genes with the largest absolute fold changes were included in the analysis. Gene lists were submitted using official mouse gene symbols, with *Mus musculus* selected as both the input and analysis species.

Enrichment analysis was conducted against the GO Biological Process ontology using the default Metascape parameters. Enriched terms were identified using a cumulative hypergeometric test, with a minimum overlap of three genes, an enrichment factor greater than 1.5, and a nominal *P* value below 0.01. Multiple-testing-adjusted *P* values were calculated using the Benjamini–Hochberg method. Related enriched terms were grouped according to gene-membership similarity, and the most statistically significant term was selected to represent each cluster.

### Public single-cell RNA-sequencing data analysis

Publicly available single-cell RNA-sequencing datasets were analyzed in R using Seurat v5.0.1 to examine the expression of selected genes and gene signatures in intestinal epithelial cell populations.

For the Haber et al. Dataset (Haber et al., 2017), the UMI count matrix (regional_cell_sampling_UMIcounts.txt) was imported into R and used to generate a Seurat object. Gene symbols were assigned as feature names, and cell-type annotations were extracted from the cell barcodes and added to the object metadata. The count matrix was normalized using Seurat’s NormalizeData function. Expression of selected genes across the annotated cell types were visualized using Seurat DotPlot, in which dot size represents the percentage of cells expressing each gene and dot color represents the scaled average expression.

To further investigate specific tuft subpopulation the data above was re-analyzed using Seurat. Data were normalized using SCTransform, followed by principal component analysis (RunPCA), neighborhood graph construction (FindNeighbors), clustering (FindClusters), and UMAP dimensionality reduction (RunUMAP) using the first 30 principal components. Tuft cells were selected for further analysis and reclustered where appropriate. Cluster marker genes (FindAllMarkers), canonical tuft-cell markers, UCell signature scores calculated from predefined tuft-cell gene sets, and cell-cycle scores (CellCycleScoring) were jointly used to annotate tuft-cell subpopulations as Tuft 1, Tuft 2, or tuft progenitor cells. Expression of selected genes across the annotated tuft-cell populations was visualized using Seurat DotPlot, with dot size indicating the percentage of expressing cells and dot color indicating the average scaled expression.

For the Hickey et al. Dataset (Hickey et al., 2023), previously generated Seurat objects corresponding to the duodenum, ileum, and jejunum were loaded into R. Expression of established cell-type markers was first examined across all annotated cells. Tuft cells were then extracted from each regional object and merged into a single Seurat object. For regional comparisons, proximal and mid-jejunum annotations were combined under a single Jejunum label. Expression of selected genes was visualized using Seurat DotPlot, showing the scaled average expression and the percentage of cells expressing each gene. Gene-signature scores were calculated using Seurat’s AddModuleScore function.

## Supplementary tables

**<u>Table S1</u>**

Bulk RNA-seq of CD45^WT^ and CD45^ΔIEC^ tuft cells and EpCAM+ IECs.

**<u>Table S2</u>**

Bulk RNA-seq of *Hp* infected CD45^WT^ and CD45^ΔIEC^ tuft cells.

**<u>Table S3</u>**

Bulk RNA-seq of CD45^WT^ and CD45^ΔIEC^ organoids, non-treated (NT) or treated with IL-13 for 72h.

**<u>Table S4</u>**

Tuft activation gene signature.

**<u>Table S5</u>**

Low-input LC-MS proteins identified in tuft cells isolated from CD45^WT^ and CD45^ΔIEC^ mice.

**<u>Table S6</u>**

Genotyping primers.

**<u>Table S7</u>**

qPCR primers.

## Supplementary Figure Legends

**Figure S1.**
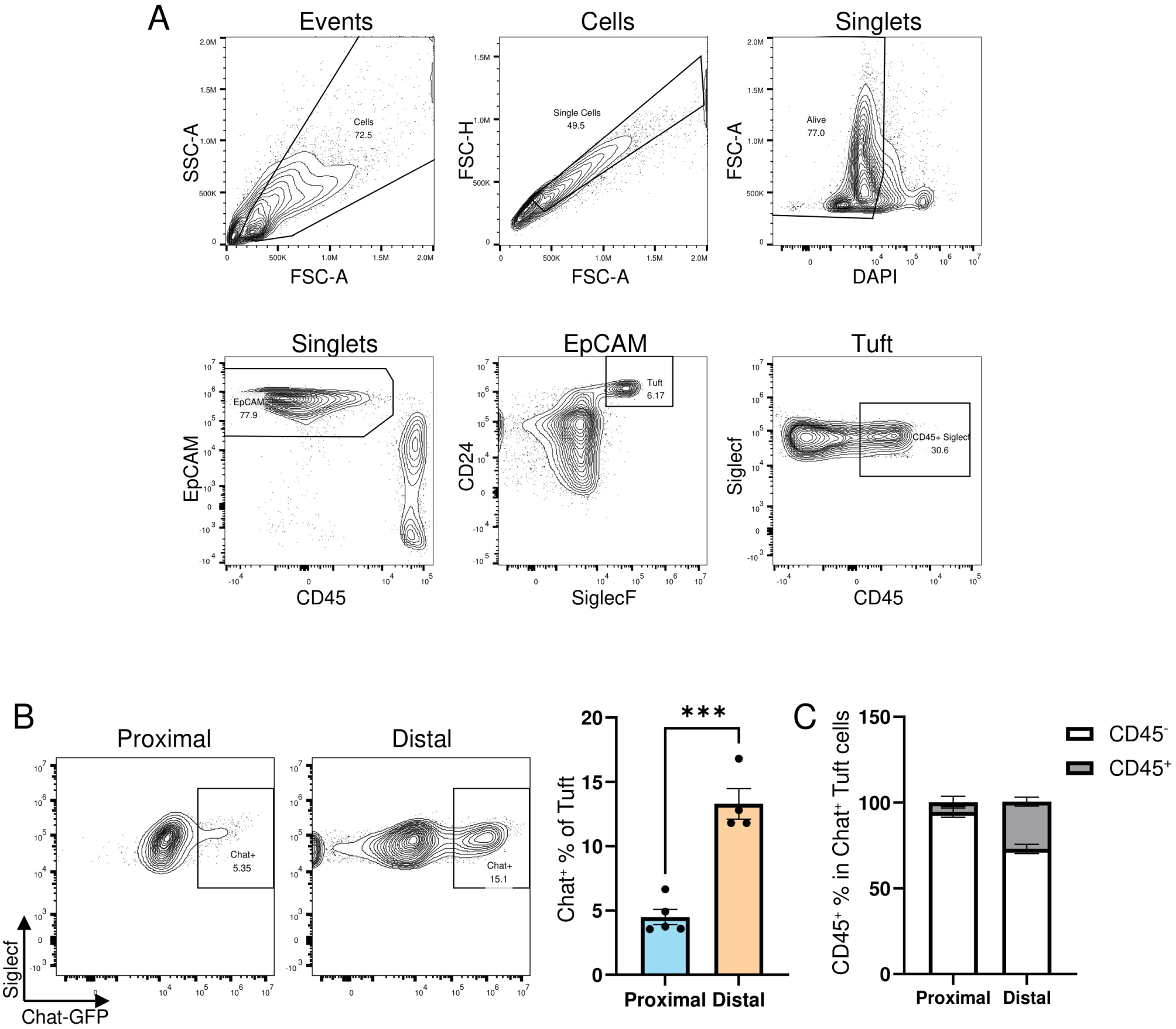
Identification of CD45-expressing tuft cells in the proximal and distal small intestine. **(A)** Representative flow cytometry gating strategy for identification of CD45⁺ tuft cells. **(B)** Representative flow cytometry plots and quantification of Chat-GFP⁺ tuft cells in the proximal and distal small intestine. n = 5 mice **(C)** Quantification of CD45⁺ and CD45^-^ tuft cells within the Chat-GFP⁺ tuft-cell population in the different regions. Data are shown as mean ± SEM. Statistical significance was determined using an unpaired two-tailed Student’s *t*-test. \*\*\**P* ≤ 0.001.

**Figure S2.**
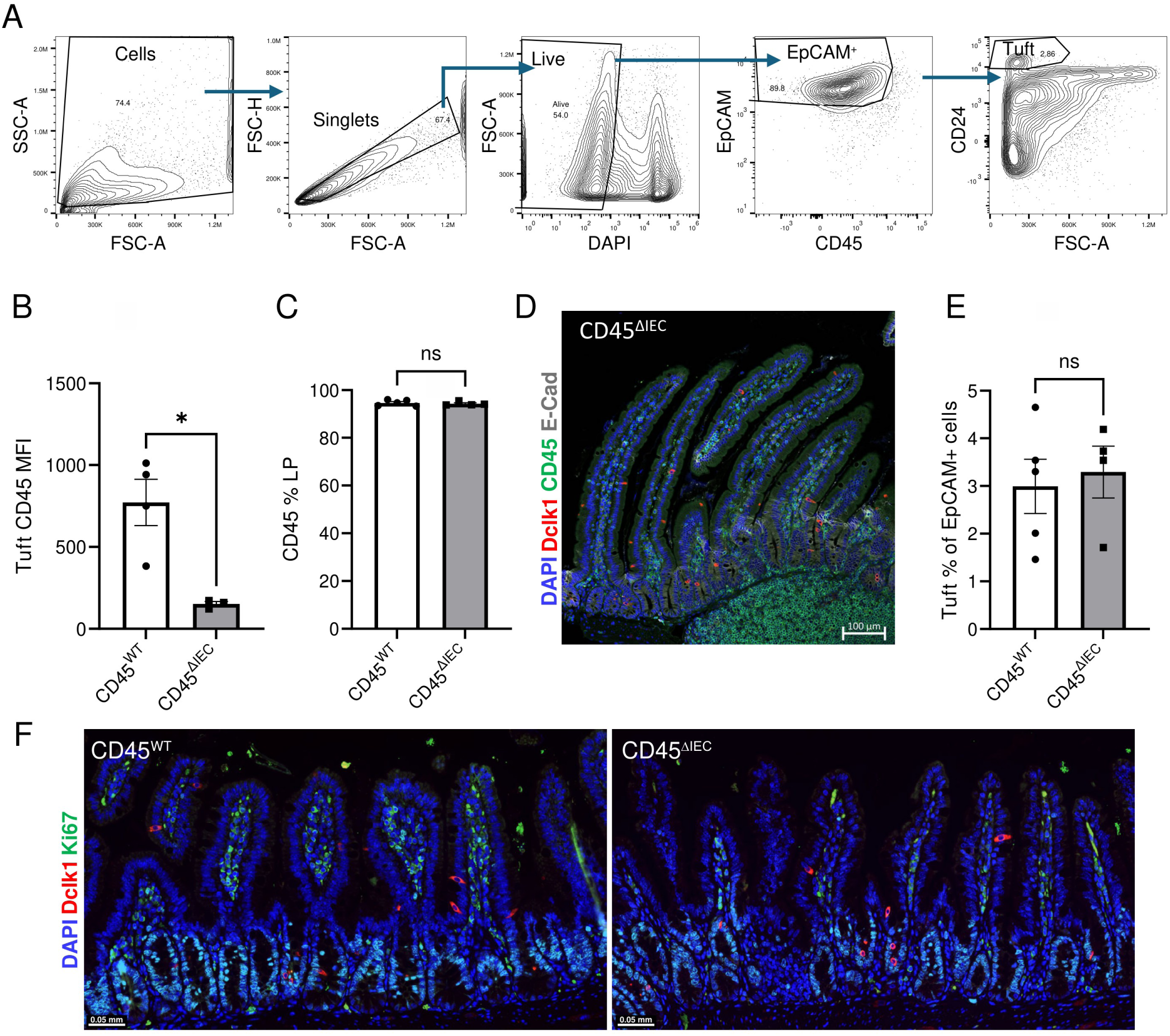
Validation of epithelial-specific CD45 deletion. **(A)** Flow cytometry gating strategy used for tuft-cell isolation. **(B)** Quantification of CD45 mean fluorescence intensity (MFI) in tuft cells from CD45^ΔIEC^ and CD45^WT^ mice. *n* = 3-4 mice per group. **(C)** CD45 expression in CD4⁺ T cells from CD45^ΔIEC^ and CD45^WT^ mice. *n* = 4-5 mice per group. **(D)** Representative immunofluorescence image stained for DCLK1 (red), CD45 (green), E-cadherin (gray), and DAPI (blue), showing preserved CD45 expression in immune cells of Peyer’s patches from CD45^ΔIEC^ mice. Scale bar 100μm. **(E)** Quantification of tuft cell frequency in CD45^ΔIEC^ and CD45^WT^ mice. *n* = 4-5 mice per group. **(F)** Representative immunofluorescence images of the distal small intestine stained for DCLK1 (red), Ki67 (green), and DAPI (blue). Data are shown as mean ± SEM. Statistical significance was determined using an unpaired two-tailed Student’s *t*-test.

**Figure S3.**
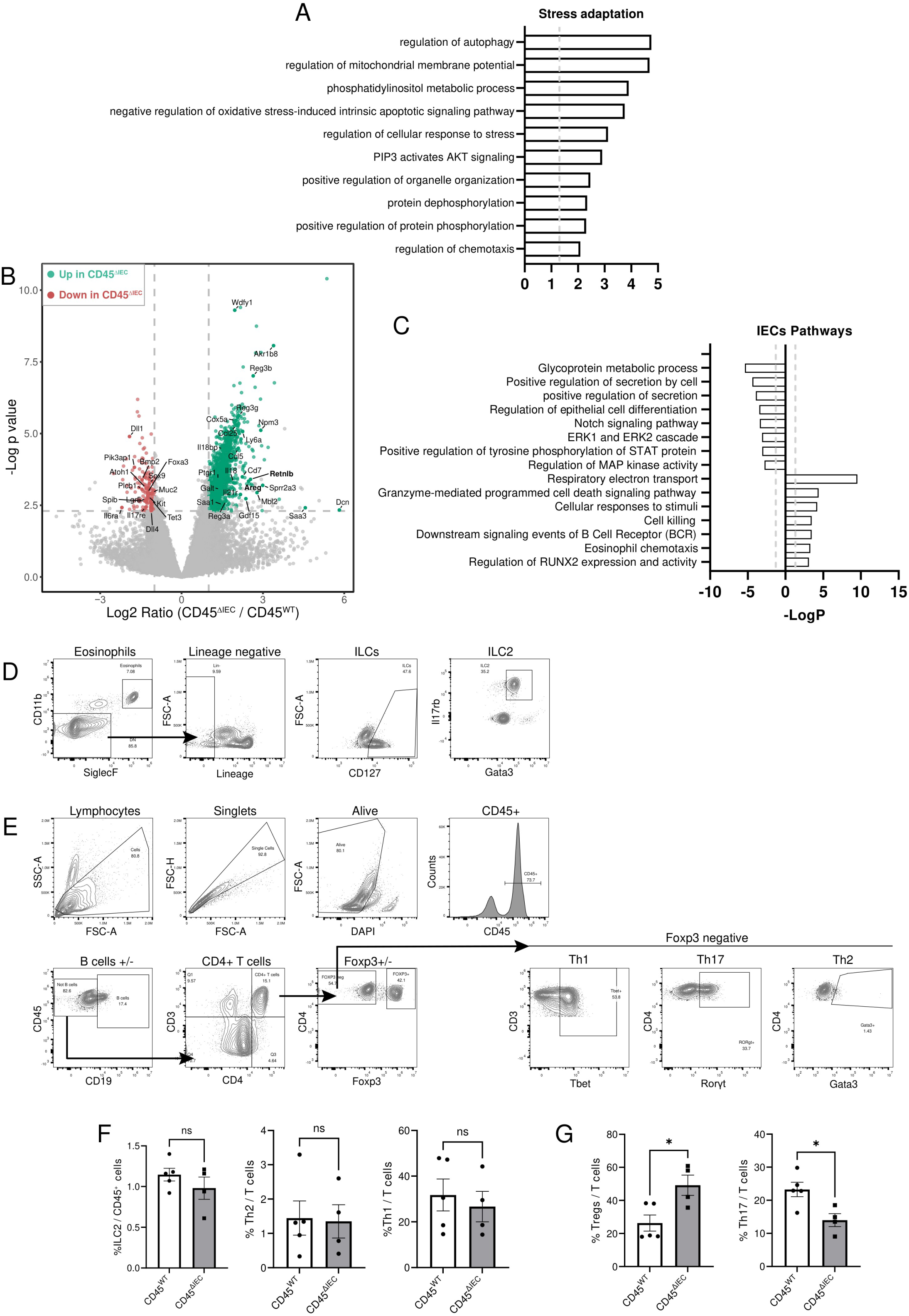
Epithelial and immune alterations following epithelial CD45 deletion. **(A)** Gene Ontology enrichment analysis of genes upregulated in CD45^ΔIEC^ tuft cells. **(B)** Volcano plot showing differentially expressed genes in intestinal epithelial cells (IECs) isolated from CD45^ΔIEC^ and CD45^WT^ mice. Selected genes are annotated. **(C)** Gene Ontology enrichment analysis of genes upregulated or downregulated in IECs. **(D)** Representative flow cytometry gating strategy for lamina propria eosinophils and ILC2s. **(E)** Representative flow cytometry gating strategy for lamina propria CD4⁺ T cell subsets. **(F)** Quantification of lamina propria ILC2, Th2, and Th1 cells from CD45^ΔIEC^ and CD45^WT^ mice. *n* = 4-5 mice per group. **(G)** Quantification of lamina propria regulatory T cells (Tregs) and Th17 cells. *n* = 4-5 mice per group. Data are shown as mean ± SEM. Statistical significance was determined using an unpaired two-tailed Student’s *t*-test. \**P* ≤ 0.05; ns, not significant.

**Figure S4.**
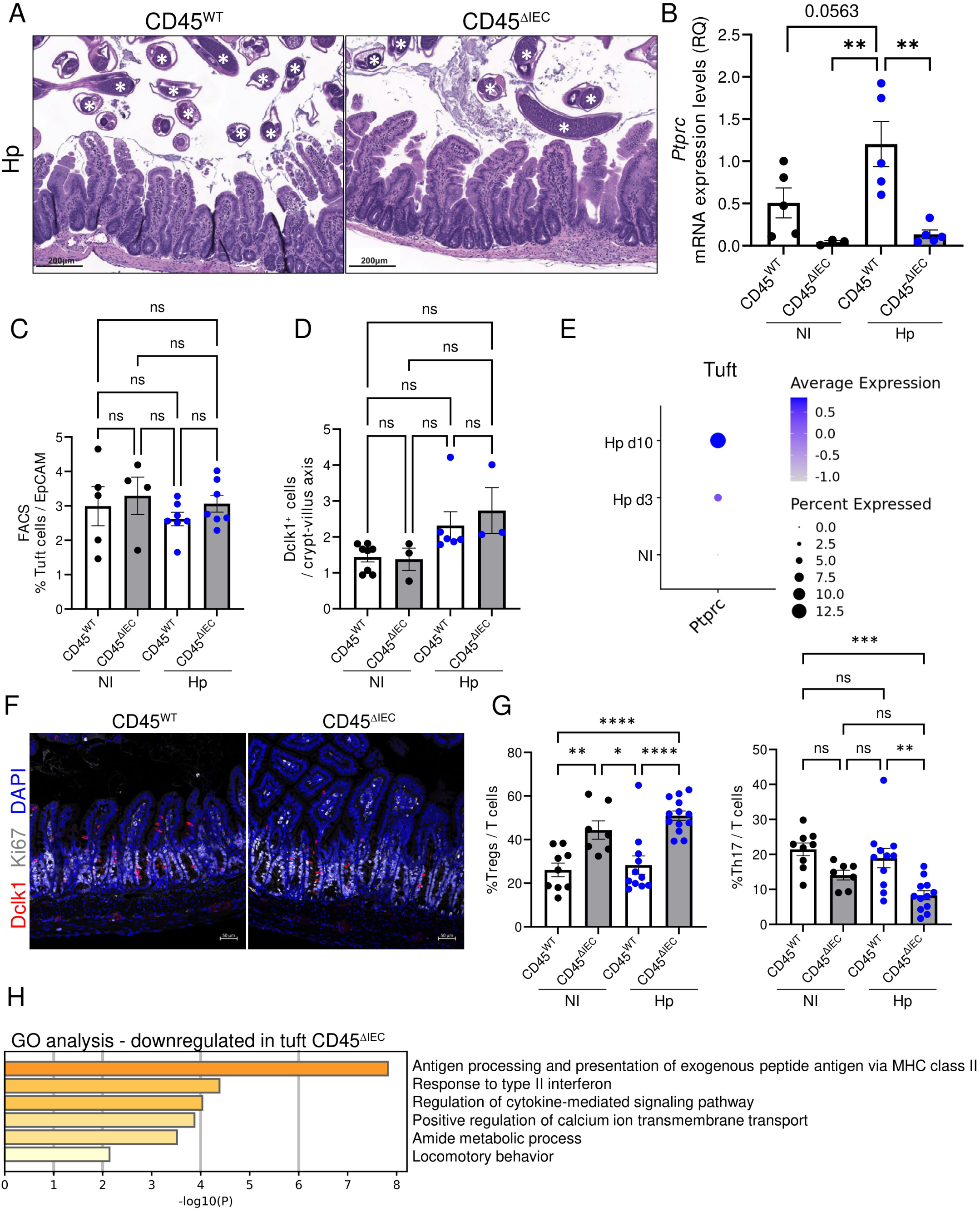
Characterization of CD45^ΔIEC^ mice during *Heligmosomoides polygyrus* infection. **(A)** Representative hematoxylin and eosin (H&E)-stained sections of proximal small intestine from *Hp*-infected CD45^ΔIEC^ and CD45^WT^ mice, Asterisks mark *Hp* worms **(B)** qPCR analysis of *Ptprc* expression in tuft cells isolated from non-infected (NI) and *Hp*-infected CD45^ΔIEC^ and CD45^WT^ mice. *n* = 3-5 mice per group. **(C)** Quantification of tuft cell frequency in the distal small intestine by flow cytometry from CD45^ΔIEC^ and CD45^WT^ non-infected (NI) or *Hp*-infected mice. *n* = 4-7 mice per group. **(D)** Quantification of DCLK1⁺ tuft cells along the crypt-villus axis by immunofluorescence analysis from CD45^ΔIEC^ and CD45^WT^ non-infected (NI) or *Hp*-infected mice. *n* = 3-8 mice per group. **(E)** Re-analysis of published single-cell RNA sequencing data (Haber et al., 2017) showing a dot plot of *Ptprc* expression within tuft cells from non-infected or infected with *Hp* for 3 or 10 days. **(F)** Representative immunofluorescence images of the distal small intestine from *Hp*-infected mice stained for DCLK1 (red), Ki67 (white), and DAPI (blue). **(G)** Quantification of lamina propria Tregs and Th17 cells out of CD4^+^ T cells by flow cytometry from CD45^ΔIEC^ and CD45^WT^ non-infected (NI) or *Hp*-infected mice. *n* = 7-13 mice per group. **(H)** Gene Ontology enrichment analysis of genes downregulated in tuft cells isolated from *Hp*-infected CD45^ΔIEC^ and CD45^WT^ mice. Data are shown as mean ± SEM. Statistical significance was determined using one-way ANOVA followed by Tukey’s multiple-comparison test. \**P* ≤ 0.05; \*\**P* ≤ 0.01; \*\*\**P* ≤ 0.001; \*\*\*\**P* ≤ 0.0001; ns, not significant.

**Figure S5.**
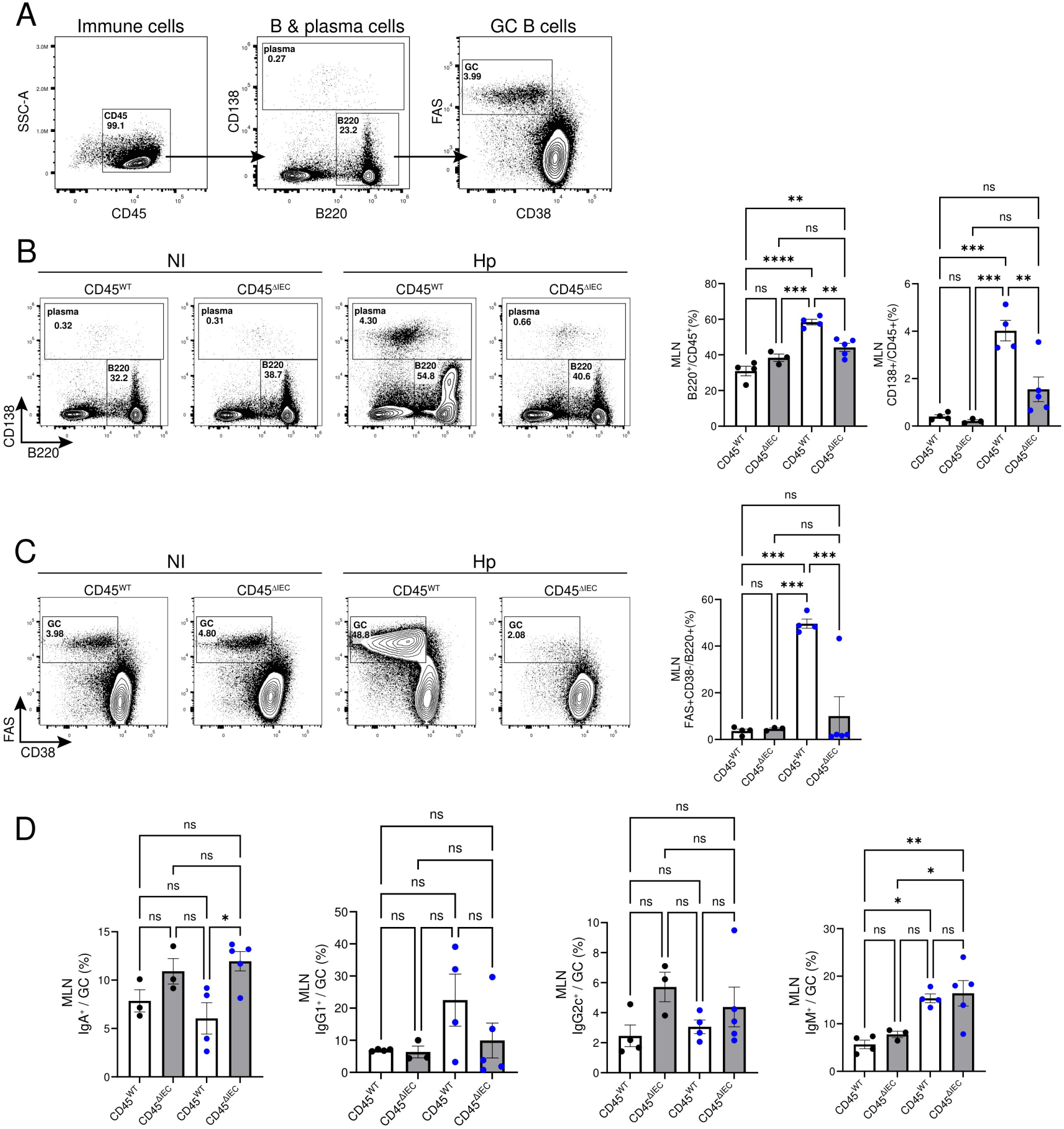
Mesenteric lymph node immune responses following *Heligmosomoides polygyrus* infection. **(A)** Representative flow cytometry gating strategy for B cell populations in mesenteric lymph nodes (MLNs). B cells (CD45^+^ B220^+^), plasma cells (CD45^+^ B220^-^ CD138^+^) and GC B cells (CD45^+^ B220^+^ CD38^-^ FAS^+^). **(B)** Representative flow cytometry plots (left) and quantification of B cells (mid) and plasma cells (right) isolated from *Hp*-infected or non-infected (NI) CD45^ΔIEC^ and CD45^WT^ MLNs. **(C)** Representative flow cytometry plots (left) and quantification of germinal-center B cells (right) isolated from *Hp*-infected or non-infected (NI) CD45^ΔIEC^ and CD45^WT^ MLNs. **(D)** Quantification of IgA⁺, IgG1⁺, IgG2c⁺, and IgM⁺ B-cell populations from *Hp*-infected or non-infected (NI) CD45^ΔIEC^ and CD45^WT^ mice. *n* = 3-5 mice per group. Data are shown as mean ± SEM. Statistical significance was determined using two-way ANOVA followed by Tukey’s multiple-comparison test. \**P* ≤ 0.05; \*\**P* ≤ 0.01; \*\*\**P* ≤ 0.001; \*\*\*\**P* ≤ 0.0001; ns, not significant.

**Figure S6.**
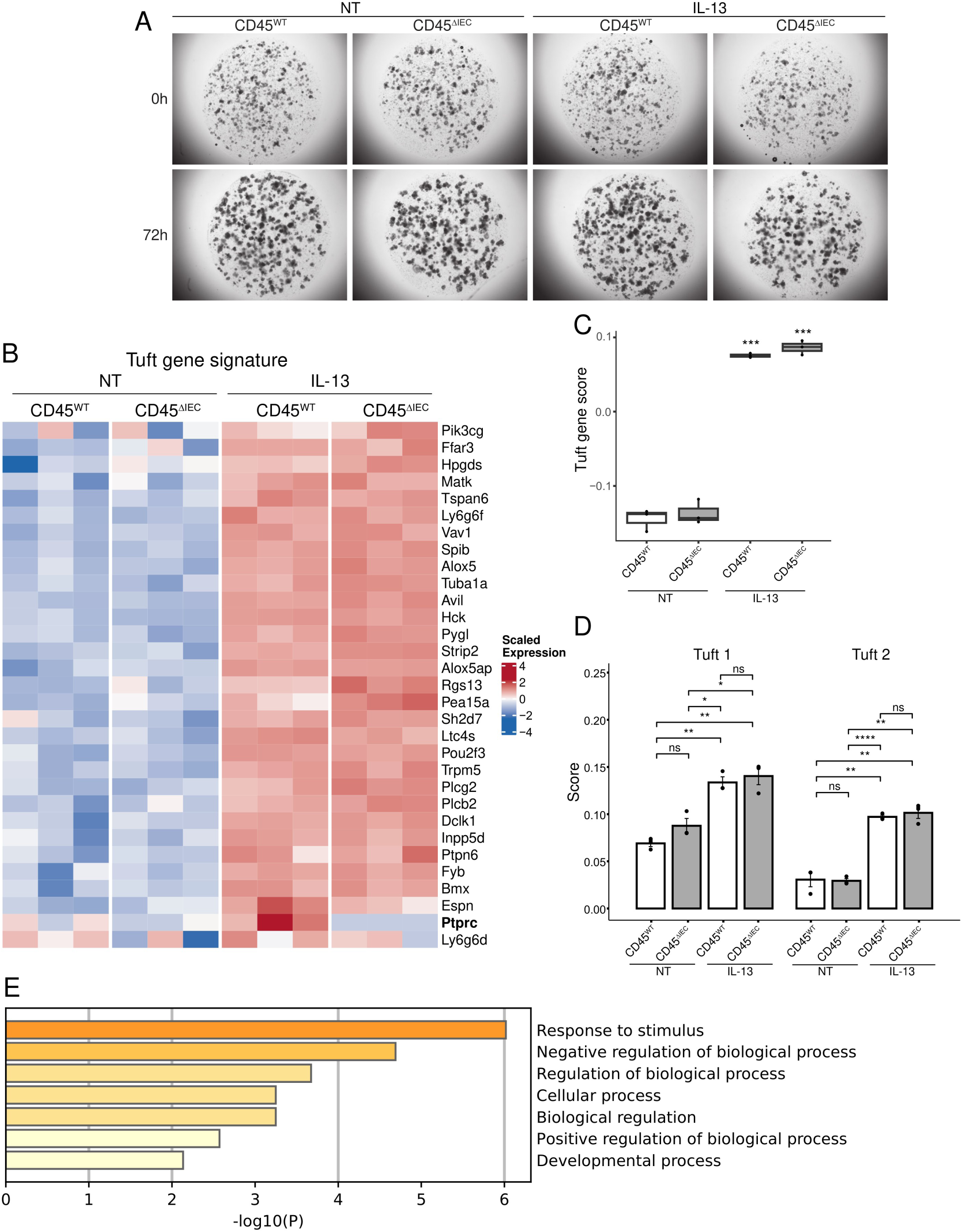
CD45 does not alter IL-13-induced tuft-cell differentiation. **(A)** Representative bright-field images of non-treated (NT) and IL-13-treated organoids at 0 and 72 h. **(B)** Heatmap showing the expression of curated tuft-cell signature genes from non-treated (NT) and IL-13-treated CD45^ΔIEC^ and CD45^WT^ organoids. *Ptprc* highlighted in bold. **(C-D)** Tuft cell subset signature scores from Haber et al. (Haber et al., 2017) in non-treated (NT) and IL-13-treated CD45^ΔIEC^ and CD45^WT^ organoids. **(C)** Tuft cell score **(D)** Tuft-1 and tuft-2 scores. **(E)** Gene Ontology enrichment analysis of the 40-gene CD45-restrained tuft-cell activation signature. *n* = 3 replicates per group. Data are shown as mean ± SEM. Statistical significance was determined using one-way ANOVA followed by Tukey’s multiple-comparison test. \**P* ≤ 0.05; \*\**P* ≤ 0.01; \*\*\**P* ≤ 0.001; \*\*\*\**P* ≤ 0.0001; ns, not significant.

**Figure S7.**
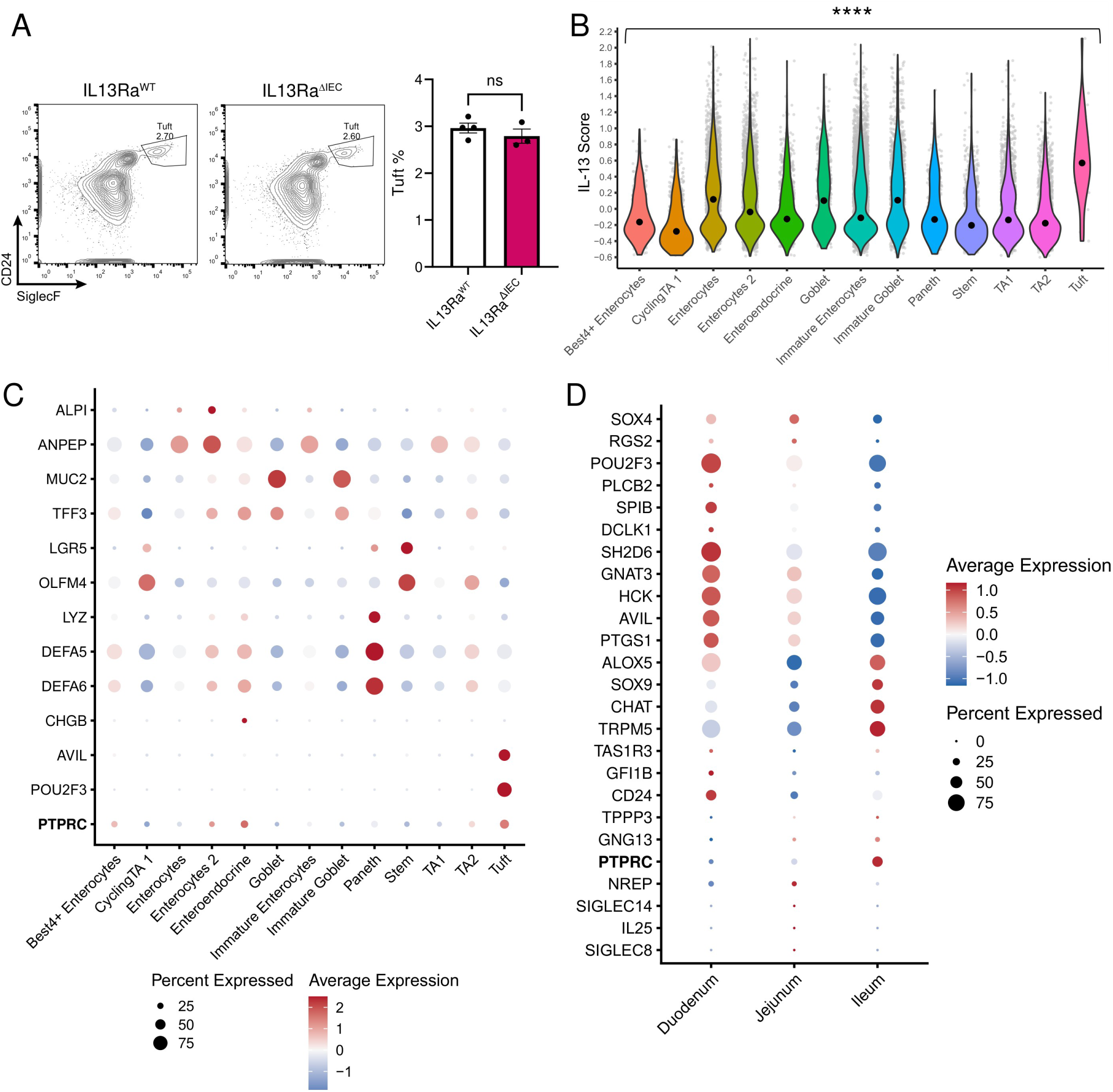
Conservation of IL-13 signaling and CD45 expression in human tuft cells. **(A)** Representative flow cytometry plots (left) and quantification of tuft cells (right) from the distal small intestine of Il-13ra1^fl/fl^ and Il-13ra1^ΔIEC^ mice. *n* = 3-4 mice per group. Data are shown as mean ± SEM. Statistical significance was determined using an unpaired two-tailed Student’s *t*-test; ns, not significant. **(B-D)** Re-analysis of published human small intestinal single-cell RNA sequencing datasets (Hickey et al., 2023). **(B)** IL-13 signaling pathway signature score (*IL13RA, IL13RB, IL4RA, STAT6*, **Methods**) across human IEC populations. \*\*\*\**P* < 0.0001 **(C)** Dot plot showing epithelial cell subset marker genes, including *PTPRC*. **(D)** Dot plot of regional expression of tuft cell signature genes, including *PTPRC*.

**Figure S8.**
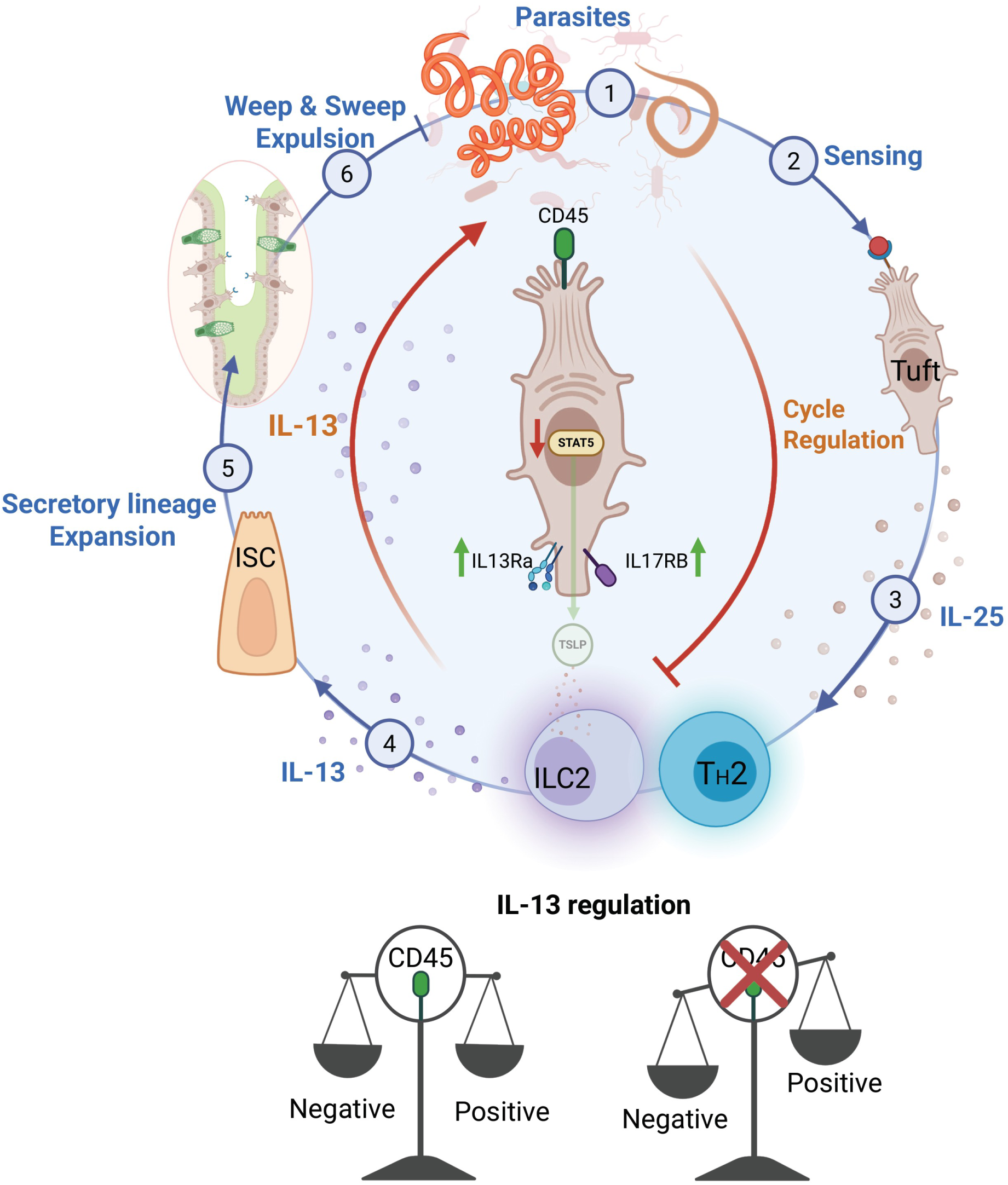
Proposed model for CD45-mediated intrinsic regulation of tuft cell type 2 immunity. Model illustrating the canonical tuft cell-ILC2/Th2 feed-forward circuit and the CD45-dependent negative-feedback pathway identified in this study. During *Heligmosomoides polygyrus* infection, tuft cells initiate type 2 immunity by producing IL-25, leading to ILC2 and Th2 activation and subsequently IL-13 secretion. IL-13, in turn, promotes tuft cell expansion while simultaneously inducing CD45 expression in activated tuft cells. CD45 restrains tuft cell activation, potentially by altering IL17RB and STAT5A protein expression, thereby limiting amplification of the type 2 immune response. Schematics were created with BioRender.

## References

Al Barashdi, M.A., A. Ali, M.F. McMullin, and K. Mills. 2021. Protein tyrosine phosphatase receptor type C (PTPRC or CD45). J. Clin. Pathol. 74:548–552.

Artis, D., M.L. Wang, S.A. Keilbaugh, W. He, M. Brenes, G.P. Swain, P.A. Knight, D.D. Donaldson, M.A. Lazar, H.R.P. Miller, G.A. Schad, P. Scott, and G.D. Wu. 2004. RELMbeta/FIZZ2 is a goblet cell-specific immune-effector molecule in the gastrointestinal tract. Proc. Natl. Acad. Sci. U. S. A. 101:13596–13600.

Banerjee, A., C.A. Herring, B. Chen, H. Kim, A.J. Simmons, A.N. Southard-Smith, M.M. Allaman, J.R. White, M.C. Macedonia, E.T. Mckinley, M.A. Ramirez-Solano, E.A. Scoville, Q. Liu, K.T. Wilson, R.J. Coffey, M.K. Washington, J.A. Goettel, and K.S. Lau. 2020. Succinate produced by intestinal microbes promotes specification of tuft cells to suppress ileal inflammation. Gastroenterology. 159:2101–2115.e5.

Belkaid, Y., and D. Artis. 2013. Immunity at the barriers. Eur. J. Immunol. 43:3096–3097.

Billipp, T.E., C. Fung, L.M. Webeck, D.B. Sargent, M.B. Gologorsky, Z. Chen, M.M. McDaniel, D.N. Kasal, J.W. McGinty, K.A. Barrow, L.M. Rich, A. Barilli, M. Sabat, J.S. Debley, C. Wu, R. Myers, M.R. Howitt, and J. von Moltke. 2024. Tuft cell-derived acetylcholine promotes epithelial chloride secretion and intestinal helminth clearance. Immunity. 57:1243–1259.e8.

Biton, M., A.L. Haber, N. Rogel, G. Burgin, S. Beyaz, A. Schnell, O. Ashenberg, C.-W. Su, C. Smillie, K. Shekhar, Z. Chen, C. Wu, J. Ordovas-Montanes, D. Alvarez, R.H. Herbst, M. Zhang, I. Tirosh, D. Dionne, L.T. Nguyen, M.E. Xifaras, A.K. Shalek, U.H. Von Andrian, D.B. Graham, O. Rozenblatt-Rosen, H.N. Shi, V. Kuchroo, O.H. Yilmaz, A. Regev, and R.J. Xavier. 2018. T Helper Cell Cytokines Modulate Intestinal Stem Cell Renewal and Differentiation. Cell. 175:1307–1320.e22.

Buissant des Amorie, J.R., M.A. Betjes, J.H. Bernink, J.H. Hageman, V.E. Geurts, H. Begthel, D. Laskaris, M.C. Heinz, I. Jordens, T. Vinck, R.M. Houtekamer, I. Verlaan-Klink, S.R. Brunner, J. van Rheenen, M. Gloerich, H. Clevers, S.J. Tans, J.S. van Zon, and H.J.G. Snippert. 2025. Intestinal tuft cell subtypes represent successive stages of maturation driven by crypt-villus signaling gradients. Nat. Commun. 16:6765.

Chen, J., R. He, W. Sun, R. Gao, Q. Peng, L. Zhu, Y. Du, X. Ma, X. Guo, H. Zhang, C. Tan, J. Wang, W. Zhang, X. Weng, J. Man, H. Bauer, Q.K. Wang, B.N. Martin, C.-J. Zhang, X. Li, and C. Wang. 2020. TAGAP instructs Th17 differentiation by bridging Dectin activation to EPHB2 signaling in innate antifungal response. Nat. Commun. 11:1913.

Chen, Y., R. Yang, B. Qi, and Z. Shan. 2024. Peptidoglycan-Chi3l1 interaction shapes gut microbiota in intestinal mucus layer. Elife. 13. doi:10.7554/eLife.92994.

Churchill, M.J., A. Pandeya, R. Bauer, T. Christopher, S. Krug, R. Honodel, S. Smita, L. Warner, B.M. Mooney, A.R. Gibson, P.S. Mitchell, E.D. Tait Wojno, and I. Rauch. 2025. Enteric tuft cell inflammasome activation drives NKp46+ILC3 IL22 via PGD2 and inhibits Salmonella. J. Exp. Med. 222:e20230803.

Concordet, J.-P., and M. Haeussler. 2018. CRISPOR: intuitive guide selection for CRISPR/Cas9 genome editing experiments and screens. Nucleic Acids Res. 46:W242–W245.

Courtney, A.H., A.A. Shvets, W. Lu, G. Griffante, M. Mollenauer, V. Horkova, W.-L. Lo, S. Yu, O. Stepanek, A.K. Chakraborty, and A. Weiss. 2019. CD45 functions as a signaling gatekeeper in T cells. Sci. Signal. 12:eaaw8151.

Delbaere, K., I. Roegiers, A. Bron, C. Durif, T. Van de Wiele, S. Blanquet-Diot, and L. Marinelli. 2023. The small intestine: dining table of host-microbiota meetings. FEMS Microbiol. Rev. 47:fuad022.

Demichev, V., C.B. Messner, S.I. Vernardis, K.S. Lilley, and M. Ralser. 2020. DIA-NN: neural networks and interference correction enable deep proteome coverage in high throughput. Nat. Methods. 17:41–44.

Ding, Z., T. Cai, J. Tang, H. Sun, X. Qi, Y. Zhang, Y. Ji, L. Yuan, H. Chang, Y. Ma, H. Zhou, L. Li, H. Sheng, and J. Qiu. 2022. Setd2 supports GATA3+ST2+ thymic-derived Treg cells and suppresses intestinal inflammation. Nat. Commun. 13:7468.

Doench, J.G., N. Fusi, M. Sullender, M. Hegde, E.W. Vaimberg, K.F. Donovan, I. Smith, Z. Tothova, C. Wilen, R. Orchard, H.W. Virgin, J. Listgarten, and D.E. Root. 2016. Optimized sgRNA design to maximize activity and minimize off-target effects of CRISPR-Cas9. Nat. Biotechnol. 34:184–191.

Earley, Z.M., W. Lisicka, J.J. Sifakis, R. Aguirre-Gamboa, A. Kowalczyk, J.T. Barlow, D.G. Shaw, V. Discepolo, I.L. Tan, S. Gona, J.D. Ernest, P. Matzinger, L.B. Barreiro, A. Morgun, A. Bendelac, R.F. Ismagilov, N. Shulzhenko, S.J. Riesenfeld, and B. Jabri. 2023. GATA4 controls regionalization of tissue immunity and commensal-driven immunopathology. Immunity. 56:43–57.e10.

Feng, X., T. Andersson, P. Flüchter, J. Gschwend, I. Berest, J.L. Muff, A. Lechner, A. Gondrand, P. Westermann, N. Brander, D. Carchidi, J.C. De Tenorio, T. Pan, U. Boehm, C.S.N. Klose, D. Artis, C.B. Messner, T. Leinders-Zufall, F. Zufall, and C. Schneider. 2025. Tuft cell IL-17RB restrains IL-25 bioavailability and reveals context-dependent ILC2 hypoproliferation. Nat. Immunol. 26:567–581.

Feng, X., P. Flüchter, J.C. De Tenorio, and C. Schneider. 2024. Tuft cells in the intestine, immunity and beyond. Nat. Rev. Gastroenterol. Hepatol. 21:852–868.

Fung, C., L.M. Fraser, G.M. Barrón, M.B. Gologorsky, S.N. Atkinson, E.R. Gerrick, M. Hayward, J. Ziegelbauer, J.A. Li, K.F. Nico, M.D.W. Tyner, L.B. DeSchepper, A. Pan, N.H. Salzman, and M.R. Howitt. 2023. Tuft cells mediate commensal remodeling of the small intestinal antimicrobial landscape. Proc. Natl. Acad. Sci. U. S. A. 120:e2216908120.

Gerbe, F., E. Sidot, D.J. Smyth, M. Ohmoto, I. Matsumoto, V. Dardalhon, P. Cesses, L. Garnier, M. Pouzolles, B. Brulin, M. Bruschi, Y. Harcus, V.S. Zimmermann, N. Taylor, R.M. Maizels, and P. Jay. 2016. Intestinal epithelial tuft cells initiate type 2 mucosal immunity to helminth parasites. Nature. 529:226–230.

Gieseck, R.L., 3rd, M.S. Wilson, and T.A. Wynn. 2018. Type 2 immunity in tissue repair and fibrosis. Nat. Rev. Immunol. 18:62–76.

Grencis, R.K. 2015. Immunity to helminths: resistance, regulation, and susceptibility to gastrointestinal nematodes. Annu. Rev. Immunol. 33:201–225.

Haber, A.L., M. Biton, N. Rogel, R.H. Herbst, K. Shekhar, C. Smillie, G. Burgin, T.M. Delorey, M.R. Howitt, Y. Katz, I. Tirosh, S. Beyaz, D. Dionne, M. Zhang, R. Raychowdhury, W.S. Garrett, O. Rozenblatt-Rosen, H.N. Shi, O. Yilmaz, R.J. Xavier, and A. Regev. 2017. A single-cell survey of the small intestinal epithelium. Nature. 551:333–339.

Harris, N., and W.C. Gause. 2011. To B or not to B: B cells and the Th2-type immune response to helminths. Trends Immunol. 32:80–88.

Haveri, H., M. Ashorn, S. Iltanen, D.B. Wilson, L.C. Andersson, and M. Heikinheimo. 2009. Enhanced expression of transcription factor GATA-4 in inflammatory bowel disease and its possible regulation by TGF-beta1. J. Clin. Immunol. 29:444–453.

He, R., J. Chen, Z. Zhao, C. Shi, Y. Du, M. Yi, L. Feng, Q. Peng, Z. Cui, R. Gao, H. Wang, Y. Huang, Z. Liu, and C. Wang. 2022. T-cell activation Rho GTPase-activating protein maintains intestinal homeostasis by regulating intestinal T helper cells differentiation through the gut microbiota. Front. Microbiol. 13:1030947.

Hennig, B.P., L. Velten, I. Racke, C.S. Tu, M. Thoms, V. Rybin, H. Besir, K. Remans, and L.M. Steinmetz. 2018. Large-scale low-cost NGS library preparation using a robust Tn5 purification and tagmentation protocol. G3 (Bethesda). 8:79–89.

Herbert, D.R., J.-Q. Yang, S.P. Hogan, K. Groschwitz, M. Khodoun, A. Munitz, T. Orekov, C. Perkins, Q. Wang, F. Brombacher, J.F. Urban Jr, M.E. Rothenberg, and F.D. Finkelman. 2009. Intestinal epithelial cell secretion of RELM-beta protects against gastrointestinal worm infection. J. Exp. Med. 206:2947–2957.

Hermiston, M.L., Z. Xu, and A. Weiss. 2003. CD45: a critical regulator of signaling thresholds in immune cells. Annu. Rev. Immunol. 21:107–137.

Hickey, J.W., W.R. Becker, S.A. Nevins, A. Horning, A.E. Perez, C. Zhu, B. Zhu, B. Wei, R. Chiu, D.C. Chen, D.L. Cotter, E.D. Esplin, A.K. Weimer, C. Caraccio, V. Venkataraaman, C.M. Schürch, S. Black, M. Brbić, K. Cao, S. Chen, W. Zhang, E. Monte, N.R. Zhang, Z. Ma, J. Leskovec, Z. Zhang, S. Lin, T. Longacre, S.K. Plevritis, Y. Lin, G.P. Nolan, W.J. Greenleaf, and M. Snyder. 2023. Organization of the human intestine at single-cell resolution. Nature. 619:572–584.

Howitt, M.R., S. Lavoie, M. Michaud, A.M. Blum, S.V. Tran, J.V. Weinstock, C.A. Gallini, K. Redding, R.F. Margolskee, L.C. Osborne, D. Artis, and W.S. Garrett. 2016. Tuft cells, taste-chemosensory cells, orchestrate parasite type 2 immunity in the gut. Science. 351:1329–1333.

Hsu, P.D., D.A. Scott, J.A. Weinstein, F.A. Ran, S. Konermann, V. Agarwala, Y. Li, E.J. Fine, X. Wu, O. Shalem, T.J. Cradick, L.A. Marraffini, G. Bao, and F. Zhang. 2013. DNA targeting specificity of RNA-guided Cas9 nucleases. Nat. Biotechnol. 31:827– 832.

Huang, L., J.H. Bernink, A. Giladi, D. Krueger, G.J.F. van Son, M.H. Geurts, G. Busslinger, L. Lin, H. Begthel, M. Zandvliet, C.J. Buskens, W.A. Bemelman, C. López-Iglesias, P.J. Peters, and H. Clevers. 2024. Tuft cells act as regenerative stem cells in the human intestine. Nature. 634:929–935.

Irie-Sasaki, J., T. Sasaki, W. Matsumoto, A. Opavsky, M. Cheng, G. Welstead, E. Griffiths, C. Krawczyk, C.D. Richardson, K. Aitken, N. Iscove, G. Koretzky, P. Johnson, P. Liu, D.M. Rothstein, and J.M. Penninger. 2001. CD45 is a JAK phosphatase and negatively regulates cytokine receptor signalling. Nature. 409:349–354.

Johnston, C.J.C., E. Robertson, Y. Harcus, J.R. Grainger, G. Coakley, D.J. Smyth, H.J. McSorley, and R. Maizels. 2015. Cultivation of Heligmosomoides polygyrus: an immunomodulatory nematode parasite and its secreted products. J. Vis. Exp. e52412.

Kumar, V., P. Cheng, T. Condamine, S. Mony, L.R. Languino, J.C. McCaffrey, N. Hockstein, M. Guarino, G. Masters, E. Penman, F. Denstman, X. Xu, D.C. Altieri, H. Du, C. Yan, and D.I. Gabrilovich. 2016. CD45 phosphatase inhibits STAT3 transcription factor activity in myeloid cells and promotes tumor-associated macrophage differentiation. Immunity. 44:303–315.

Lei, W., W. Ren, M. Ohmoto, J.F. Urban Jr, I. Matsumoto, R.F. Margolskee, and P. Jiang. 5 2018. Activation of intestinal tuft cell-expressed Sucnr1 triggers type 2 immunity in the mouse small intestine. Proc. Natl. Acad. Sci. U. S. A. 115:5552–5557.

Lindholm, H.T., N. Parmar, C. Drurey, M. Campillo Poveda, P.M. Vornewald, J. Ostrop, A. Díez-Sanchez, R.M. Maizels, and M.J. Oudhoff. 2022. BMP signaling in the intestinal epithelium drives a critical feedback loop to restrain IL-13-driven tuft cell hyperplasia. Sci. Immunol. 7:eabl6543.

Lloyd, C.M., and R.J. Snelgrove. 2018. Type 2 immunity: Expanding our view. Sci. Immunol. 3:eaat1604.

Loh, W., and M.L.K. Tang. 2018. The epidemiology of food allergy in the global context. Int. J. Environ. Res. Public Health. 15:2043.

Manco, R., I. Averbukh, Z. Porat, K. Bahar Halpern, I. Amit, and S. Itzkovitz. 2021. Clump sequencing exposes the spatial expression programs of intestinal secretory cells. Nat. Commun. 12:3074.

McGinty, J.W., H.-A. Ting, T.E. Billipp, M.S. Nadjsombati, D.M. Khan, N.A. Barrett, H.-E. Liang, I. Matsumoto, and J. von Moltke. 2020. Tuft-cell-derived leukotrienes drive rapid anti-helminth immunity in the small intestine but are dispensable for anti-protist immunity. Immunity. 52:528–541.e7.

Medzhitov, R., D.S. Schneider, and M.P. Soares. 2012. Disease tolerance as a defense strategy. Science. 335:936–941.

Meier, F., A.-D. Brunner, M. Frank, A. Ha, I. Bludau, E. Voytik, S. Kaspar-Schoenefeld, M. Lubeck, O. Raether, N. Bache, R. Aebersold, B.C. Collins, H.L. Röst, and M. Mann. 2020. diaPASEF: parallel accumulation-serial fragmentation combined with data-independent acquisition. Nat. Methods. 17:1229–1236.

von Moltke, J., M. Ji, H.E. Liang, and R.M. Locksley. 2016. Tuft-cell-derived IL-25 regulates an intestinal ILC2-epithelial response circuit. Nature. 529:221–225.

Morimoto, Y., K. Hirahara, M. Kiuchi, T. Wada, T. Ichikawa, T. Kanno, M. Okano, K. Kokubo, A. Onodera, D. Sakurai, Y. Okamoto, and T. Nakayama. 2018. Amphiregulin-producing pathogenic memory T helper 2 cells instruct eosinophils to secrete osteopontin and facilitate airway fibrosis. Immunity. 49:134–150.e6.

Nadjsombati, M.S., J.W. McGinty, M.R. Lyons-Cohen, J.B. Jaffe, L. DiPeso, C. Schneider, C.N. Miller, J.L. Pollack, G.A. Nagana Gowda, M.F. Fontana, D.J. Erle, M.S. Anderson, R.M. Locksley, D. Raftery, and J. von Moltke. 2018. Detection of succinate by intestinal tuft cells triggers a type 2 innate immune circuit. Immunity. 49:33–41.e7.

Nevo, S., N. Frenkel, N. Kadouri, T. Gome, N. Rosenthal, T. Givony, A. Avin, C. Peligero Cruz, M. Kedmi, M. Lindzen, S. Ben Dor, G. Damari, Z. Porat, R. Haffner-Krausz, H. Keren-Shaul, Y. Yarden, A. Munitz, D. Leshkowitz, Y. Goldfarb, and J. Abramson. 2024. Tuft cells and fibroblasts promote thymus regeneration through ILC2-mediated type 2 immune response. Sci. Immunol. 9:eabq6930.

Ogulur, I., Y. Mitamura, D. Yazici, Y. Pat, S. Ardicli, M. Li, P. D’Avino, C. Beha, H. Babayev, B. Zhao, C. Zeyneloglu, O. Giannelli Viscardi, O. Ardicli, A. Kiykim, A. Garcia-Sanchez, J.-F. Lopez, L.-L. Shi, M. Yang, S.R. Schneider, S. Skolnick, R. Dhir, U. Radzikowska, A.J. Kulkarni, M.B. Imam, W. van de Veen, M. Sokolowska, M. Martin-Fontecha, O. Palomares, K.C. Nadeau, M. Akdis, and C.A. Akdis. 2025. Type 2 immunity in allergic diseases. Cell. Mol. Immunol. 22:211–242.

Oyesola, O.O., M.T. Shanahan, M. Kanke, B.M. Mooney, L.M. Webb, S. Smita, M.K. Matheson, P. Campioli, D. Pham, S.P. Früh, J.W. McGinty, M.J. Churchill, J.L. Cahoon, P. Sundaravaradan, B.A. Flitter, K. Mouli, M.S. Nadjsombati, E. Kamynina, S.A. Peng, R.L. Cubitt, K. Gronert, J.D. Lord, I. Rauch, J. von Moltke, P. Sethupathy, and E.D. Tait Wojno. 2021. PGD2 and CRTH2 counteract Type 2 cytokine-elicited intestinal epithelial responses during helminth infection. J. Exp. Med. 218. doi:10.1084/jem.20202178.

Picelli, S., Å.K. Björklund, O.R. Faridani, S. Sagasser, G. Winberg, and R. Sandberg. 2013. Smart-seq2 for sensitive full-length transcriptome profiling in single cells. Nat. Methods. 10:1096–1098.

Picelli, S., A.K. Björklund, B. Reinius, S. Sagasser, G. Winberg, and R. Sandberg. 2014a. Tn5 transposase and tagmentation procedures for massively scaled sequencing projects. Genome Res. 24:2033–2040.

Picelli, S., O.R. Faridani, A.K. Björklund, G. Winberg, S. Sagasser, and R. Sandberg. 2014b. Full-length RNA-seq from single cells using Smart-seq2. Nat. Protoc. 9:171–181.

Reynolds, L.A., K.J. Filbey, and R.M. Maizels. 2012. Immunity to the model intestinal helminth parasite Heligmosomoides polygyrus. Semin. Immunopathol. 34:829–846.

Roach, S.N., J.K. Fiege, F.K. Shepherd, T.D. Wiggen, R.C. Hunter, and R.A. Langlois. 2022. Respiratory Influenza virus infection causes dynamic tuft cell and innate lymphoid cell changes in the small intestine. J. Virol. 96:e0035222.

Saunders, A.E., and P. Johnson. 2010. Modulation of immune cell signalling by the leukocyte common tyrosine phosphatase, CD45. Cell. Signal. 22:339–348.

Schneider, C., C.E. O’Leary, J. von Moltke, H.-E. Liang, Q.Y. Ang, P.J. Turnbaugh, S. Radhakrishnan, M. Pellizzon, A. Ma, and R.M. Locksley. 2018. A metabolite-triggered tuft cell-ILC2 circuit drives small intestinal remodeling. Cell. 174:271–284.e14.

Stephen-Victor, E., G.A. Kuziel, M. Martinez-Blanco, B.-E. Jugder, M. Benamar, Z. Wang, Q. Chen, G.L. Lozano, A. Abdel-Gadir, Y. Cui, J. Fong, E. Saint-Denis, I. Chang, K.C. Nadeau, W. Phipatanakul, A. Zhang, F.A. Farraj, F. Holder-Niles, D. Zeve, D.T. Breault, K. Schmitz-Abe, R. Rachid, E. Crestani, S. Rakoff-Nahoum, and T.A. Chatila. 2025. RELMβ sets the threshold for microbiome-dependent oral tolerance. Nature. 638:760–768.

Strine, M.S., and C.B. Wilen. 2022. Tuft cells are key mediators of interkingdom interactions at mucosal barrier surfaces. PLoS Pathog. 18:e1010318.

Tang, M.L.K., and R.J. Mullins. 2017. Food allergy: is prevalence increasing?: Food allergy prevalence time trends. Intern. Med. J. 47:256–261.

Tchilian, E.Z., and P.C.L. Beverley. 2006. Altered CD45 expression and disease. Trends Immunol. 27:146–153.

Vuik, F., J. Dicksved, S.Y. Lam, G.M. Fuhler, L. van der Laan, A. van de Winkel, S.R. Konstantinov, M. Spaander, M.P. Peppelenbosch, L. Engstrand, and E.J. Kuipers. 2019. Composition of the mucosa-associated microbiota along the entire gastrointestinal tract of human individuals. United European Gastroenterol. J. 7:897– 907.

Wang, J., R. Shen, K. Yang, X. Guo, J. Yu, C. Wu, X.-K. Guo, and X. Hu. 2025. Tuft cells restrain intestinal type 2 immunity through the transcription factor Spi-B. Sci. Immunol. 10:eads5818.

Wang, Y., M.A. Su, and Y.Y. Wan. 2011. An essential role of the transcription factor GATA-3 for the function of regulatory T cells. Immunity. 35:337–348.

Wilen, C.B., S. Lee, L.L. Hsieh, R.C. Orchard, C. Desai, B.L. Hykes Jr, M.R. McAllaster, D.R. Balce, T. Feehley, J.R. Brestoff, C.A. Hickey, C.C. Yokoyama, Y.-T. Wang, D.A. MacDuff, D. Kreamalmayer, M.R. Howitt, J.A. Neil, K. Cadwell, P.M. Allen, S.A. Handley, M. van Lookeren Campagne, M.T. Baldridge, and H.W. Virgin. 2018. Tropism for tuft cells determines immune promotion of norovirus pathogenesis. Science. 360:204–208.

Xiong, X., C. Yang, W.-Q. He, J. Yu, Y. Xin, X. Zhang, R. Huang, H. Ma, S. Xu, Z. Li, J. Ma, L. Xu, Q. Wang, K. Ren, X.S. Wu, C.R. Vakoc, J. Zhong, G. Zhong, X. Zhu, Y. Song, H.-B. Ruan, and Q. Wang. 2022. Sirtuin 6 maintains epithelial STAT6 activity to support intestinal tuft cell development and type 2 immunity. Nat. Commun. 13:5192.

Xiong, Z., X. Zhu, J. Geng, Y. Xu, R. Wu, C. Li, D. Fan, X. Qin, Y. Du, Y. Tian, and Z. Fan. 4 2022. Intestinal Tuft-2 cells exert antimicrobial immunity via sensing bacterial metabolite N-undecanoylglycine. Immunity. 55:686–700.e7.

Xu, H., Y. Wang, W. Wang, Y.-X. Fu, J. Qiu, Y. Shi, L. Yuan, C. Dong, X. Hu, Y.-G. Chen, and X. Guo. 2025. ILC3s promote intestinal tuft cell hyperplasia and anthelmintic immunity through RANK signaling. Sci. Immunol. 10:eadn1491.

Yang, Y., Y.-H. Hu, and Y. Liu. 2020. Wdfy1 deficiency impairs Tlr3-mediated immune responses in vivo. Cell. Mol. Immunol. 17:1014–1016.

Zaiss, D.M., L. Yang, P.R. Shah, J.J. Kobie, J.F. Urban, and T.R. Mosmann. 2006. Amphiregulin, a TH2 cytokine enhancing resistance to nematodes. Science. 314:1746.

Zaiss, D.M.W., W.C. Gause, L.C. Osborne, and D. Artis. 2015. Emerging functions of amphiregulin in orchestrating immunity, inflammation, and tissue repair. Immunity. 42:216–226.

